# Variations in Structural MRI Quality Significantly Impact Commonly-Used Measures of Brain Anatomy

**DOI:** 10.1101/581876

**Authors:** Alysha Gilmore, Nicholas Buser, Jamie L. Hanson

## Abstract

Subject motion can introduce noise into neuroimaging data and result in biased estimations of brain structure. In-scanner motion can compromise data quality in a number of ways and varies widely across developmental and clinical populations. However, quantification of structural image quality is often limited to proxy or indirect measures gathered from functional scans; this may be missing true differences related to these potential artifacts. In this study, we take advantage of novel informatic tools, the CAT12 toolbox, to more directly measure image quality from T1-weighted images to understand if these measures of image quality: 1) relate to rigorous quality-control checks visually completed by human raters; 2) are associated with sociodemographic variables of interest; 3) influence regional estimates of cortical surface area, cortical thickness, and subcortical volumes from the commonly-used Freesurfer tool suite. We leverage public-access data that includes a community-based sample of children and adolescents, spanning a large age-range (*N=388; ages 5-21*). Interestingly, even after visually inspecting our data, we find image quality significantly impacts derived cortical surface area, cortical thickness, and subcortical volumes from multiple regions across the brain (∼23.4% of all areas investigated). We believe these results are important for research groups completing structural MRI studies using Freesurfer or other morphometric tools. As such, future studies should consider using measures of image quality to minimize the influence of this potential confound in group comparisons or studies focused on individual differences.

## 1. INTRODUCTION

Neuroimaging methods are increasingly common, but with these advancements, there has been a greater understanding of the potential confounds and limitations of these research techniques. One of the most common limitations of neuroimaging research is that of motion-related artifacts. This type of noise is caused by participant movement during a neuroimaging session and may impact assessment of brain structure and function [1–4]. For those interested in neurodevelopment and mental health, such noise and bias may be particularly important to address. While head motion varies considerably among individuals, children typically move more than adults and patient groups move on average more than controls [5, 6].

Multiple resting state fMRI studies have highlighted the importance of this issue, as very small differences in motion have been shown to yield significant differences in estimates of functional connectivity among healthy samples [1, 3]. In fact, head movements within fractions of a millimeter have been shown to significantly bias correlations between BOLD-activation time series’ in a distant dependent manner, leading to spurious estimates of connectivity within functional networks [3, 7]. Further, recent work has shown that head motion is consistent within individual subjects from one scanning session to the next, raising the potential for motion to confound the exploration of individual differences within the same population [8]. Particularly challenging, these differences persist even after extensive motion correction procedures [9, 10]. This has thus motivated a methodological sub-field focused on effective ways to reduce motion-related noise in resting-state and other forms of functional MRI.

While a great deal of progress has been made in quantifying and addressing the impact of head-motion in functional analyses, less attention has been given to structural MRI, such as estimates derived from T1-weighted images. It is, however, clear that head motion has been shown to compromise derived measures of volume and thickness in regions of cortical gray matter [11–14]. Such effects remain after different forms of manual and automatic correction, suggesting that in-scanner motion induces spurious effects that do not reflect a processing failure in software, rather, they reflect systematic bias (e.g., motion-induced blurring) and this may appear similar to gray matter atrophy [13]. Particularly concerning, many neuroimaging groups will visually inspect scans and include scans of “fair” or “marginal” quality. As researchers focus on different groups (e.g., children versus adolescents; clinical groups versus non-clinical groups), this potentially creates an “apples versus oranges” comparison; all scans may “*pass*” visual inspection, but one group has excellent image quality and clarity, while another has visible motion and is only above these passing thresholds. Such issues are sadly still ignored quite broadly in neuroimaging but have significant implications for potential results. For example, Ducharme and colleagues [15] probed potential non-linear trajectories of neurodevelopment during childhood and adolescence in a sample without any quality control (QC), with standard QC, and also more stringent QC. Using no QC, 16.4% of the brain showed either quadratic or cubic developmental trajectories; this however dropped to 9.7% and 1.4% of the brain for standard and more stringent quality control. Such patterns strongly underscore the importance of these issues when working with pediatric, clinical, or any other potential “high-motion” populations.

While the impact of movement on structural MRI is clear, methods of quantifying and addressing motion-related noise in T1-weighted images have been limited. With particularly noisy structural data, researchers traditionally “flag” problematic scans and remove these subjects from further analyses. This process involves raters visually assessing each T1-weighted structural image. A limitation of this strategy is that many phenotypes of interest are inherently more prone to head motion (e.g., children under 9; individuals with clinical diagnoses [12, 14]). Also, human rating systems are relatively impractical for large scale datasets. A further challenge is that visual inspection by human raters is relatively subjective. Numerous studies have showcased this, with moderately concerning inter-and intra-related variability among human-rating systems [16]. Further, even for T1-weighted scans that pass “visual inspection”, there may still be important variations in data quality which impact morphometric estimates. As noted previously and put another way, some scans may be “just above” threshold for raters, while other volumes may be of utmost quality; both types of scans, however, would be simply considered “usable” [12].

Thinking holistically, these multiple problems are in part due to the limited information about noise typically available for T1-weighted MRI scans. T1-weighted MRI scans involve the acquisition of only one, higher resolution anatomical volume. To date, this has prohibited rich assessments of noise and subject movement in contrast to fMRI. Functional MRI involves the acquisition of dozens, often hundreds, of lower resolution brain volumes; this allows for the calculation of frame-by-frame changes in a volume’s position, and a clear metric of subject movement during fMRI scanning acquisitions. The ease in collection of this sort of data has led some to advocate for the use of fMRI-derived motion parameters, such as mean Framewise Displacement (FD), to identify structural brain scans that contain motion related bias. Recent work has showed that by additionally removing FD outliers from a sample of visually inspected T1-weighted images, the effect sizes of age and gray matter thickness were attenuated across a majority of the cortex [17]. It is, therefore, possible that some past results of associations between participant variables and brain morphometry derived from T1-weighted images may be inaccurate, likely particularly inflated in “motion-prone” populations. Additional work would be necessary to clarify precisely how motion-related bias and noise in T1-weighted images varies and overlaps across distinct study populations.

While past structural MRI studies with T1-weighted images have suffered from the limitations noted above, advancements of novel informatic tools may overcome these issues. Quality assessment tools have been recently introduced that provide easy-to-implement, automated, quantitative measures of neuroimaging data. For example, the MRI Quality Control tool (MRIQC) has recently been introduced and can speak to different quality attributes of T1-weighted (and other MRI) images [18]. Similarly, the Computational Anatomy Toolbox for SPM (CAT12) assesses multiple image quality metrics and provides an aggregate “*grade*” for a given structural MRI scan [19]. Thinking about past research, it is unclear if structural MRI quality is related to commonly derived structural measures (e.g., cortical surface area; cortical thickness; regional subcortical volumes). Thoughtful work by Rosen and colleagues [20] began to investigate this idea. These researchers found that metrics from Freesurfer, specifically Euler number, were consistently correlated with human raters’ assessments of image quality. Furthermore, Euler number, a summary statistic of the topological complexity of a reconstructed brain surface, was significantly related to variations in cortical thickness.

While important, one of Rosen and colleagues’ major results could be described as “collinear” in nature--a measure of Freesurfer re-construction (Euler number) is related to measures output by Freesurfer (cortical thickness) [20]. In theory, inaccuracy or variability of Freesurfer re-construction could be due to MR quality, or algorithmic issues. The use of an independent measure of quality in relation to Freesurfer outputs would provide stronger evidence of the potential impact of T1-weighted MRI quality on morphometric measures. In addition, Rosen and colleagues did not investigate if Euler number, their measure of MR quality, was related to subcortical (e.g., amygdala) volumes or cortical surface area. Given the major interest from cognitive and affective neuroscientists in these type of morphometric measures [21, 22], it will be important to know if T1-weighted image quality impacts variations in these structures. Accounting for such variations may be important in reducing potential spurious associations and increasing the replicability of effects.

To these ends, we investigated three key questions: 1) if an integrated measure of image quality, output by the CAT12 toolbox, uniquely related to visual rater judgement (retain/exclude) of structural MRI images; 2) if variations in image quality related to sociodemographic and psychosocial variables (e.g., age; sex; clinical diagnosis); 3) if CAT12 image quality was associated with differences in commonly-used morphometric measures derived from T1-weighted images in Freesurfer (cortical surface area, cortical thickness, and subcortical volume).

## 2. MATERIALS AND METHODS

### 2.1 Participants

Data from 388 participants between the ages of 5-21 years of age with T1-weighted structural images were downloaded from two data waves of an ongoing research initiative, The Healthy Brain Network (HBN), launched by The Child Mind Institute in 2015. For sample characteristics, see Table 1. Participants with cognitive or behavioral challenges (e.g., being nonverbal, IQ<66), or with medical concerns expected to confound brain-related findings were excluded from the HBN project. The HBN protocol spans four sessions, each approximately three hours in duration. For additional information about the full HBN sample and measures, please see the HBN data-descriptor [23].

**Table 1.** Demographic Table. Table displaying demographic characteristics of our sample, including participant age, sex, psychiatric diagnosis (binary indicator based on structured interview), general cognitive ability, and body mass index. This table also displays the mean (and standard deviations) for our MRI quality metric of interest, CAT12 scores, as well as Freesurfer’s Euler Number.

| **Sample Characteristics** | **N = 388<sup>†</sup>** |
| --- | --- |
| **Sex** |  |
| Female | 142 (37%) |
| Male | 246 (63%) |
| **Age (in Years)** | 10.1 (3.4) |
| **Diagnosis** |  |
| No History | 60 (16%) |
| One or more Disorder | 307 (84%) |
| [Missing] | 21 |
| **General Cognitive Ability** | 98 (17) |
| [Missing] | 52 |
| **BMI** | 19.5 (5.2) |
| [Missing] | 8 |
| **Structural MRI Quality, CAT12 Toolbox** | 0.83 (0.07) |
| [Missing] | 1 |
| **Freesurfer Euler Number** | 113 (107) |
| [Missing] | 24 |
| <sup>†</sup> n (%); Mean (SD) |  |

### 2.2 MRI Data Acquisition

MRI acquisition included structural MRI (T1-and T2-weighted), magnetization transfer imaging, and quantitative T1-and T2-weighted mapping. Here, we focused on only T1-weighted structural MRI scans. A Siemens 3-Telsa Tim Trio MRI scanner located at the Rutgers University Brain Imaging Center (RU) was equipped with a Siemens 32-channel head coil. T1-weighted scans were acquired with a Magnetization Prepared -RApid Gradient Echo (MPRAGE) sequence with the following parameters: 224 Slices, 0.8×0.8×0.8 mm resolution, TR=2500 ms, TE=3.15 ms, and Flip Angle=8°. All neuroimaging data used in this study are openly available for download with proper data usage agreement via the International Neuroimaging Data-sharing Initiative (http://fcon_1000.projects.nitrc.org/indi/cmi_healthy_brain_network/). Again, please see the HBN data descriptor for additional information [23].

### 2.3 Visual Quality Inspection

All T1-weighted scans were separated by release wave then visually inspected by a series of human raters that were trained to recognize frequent indications of scan artifacts and motion. This training provided examples and descriptions for artifacts including “ringing”, “ghosting”, “RF-Noise”, “head coverage”, and “susceptibility”. Examples of this protocol are detailed in our *Supplemental Materials*. Each rater was instructed to give a score between 1 and 10, with high number being assigned to higher quality images. A score of a 6 was chosen as a cutoff for scan inclusion in further research. This choice was motivated by examining the mean and median of ratings from 6 research assistants who examined the structural MRI scans; the mean of all ratings was 6.14 and the median was 6. Additional information about rating distributions and correlations between rates is detailed in our Supplemental Materials. To minimize any rater idiosyncrasy, all ratings were z-scored (within rater), averaged across raters, and compared to the averaged z-score for the cutoff (6.0) points. Scans for which the averaged z-scored rating was greater than the averaged z-score cutoff point were retained (Passing N=209) and the rest were removed from further analysis. In our supplemental materials, we also completed additional analyses with subjects who did not pass <u>visual quality inspection, examining similar relations between image quality and morphometric outputs</u>.

### 2.4 Image Quality Metrics

The CAT12 toolbox (Computational Anatomy Toolbox 12) from the Structural Brain Mapping group, implemented in Statistical Parametric Mapping, was used to generate a quantitative metric indicating the quality of each T1-weighted image [19, 24]. The method employed considers four summary measures of image quality: 1) noise to contrast ratio, 2) coefficient of joint variation, 3) inhomogeneity to contrast ratio, and 4) Root mean squared voxel resolution. To produce a single aggregate metric that serves as an indicator of overall quality, this toolbox normalizes each measure and combines them using a kappa statistic-based framework, for optimizing a generalized linear model through solving least squares [25]. This measure ranged from 0-1, with higher values indicating better image quality. Additional information is available at: http://www.neuro.uni-jena.de/cat/index.html#QA. Quality assessment for one T1-weighted scan could not be completed through the CAT12 toolbox due to excessive noise. Of note, and relevant for the use of the CAT12 toolbox as a quality control tool, generation of image quality metrics took approximately 18 minutes per subject/scan (on entry-level computers, e.g., an Apple iMac with a 2.8GHz quad core Intel Core i5 processor and 16GB of RAM).

### 2.5 Sociodemographic, Cognitive, and Psychiatric Measures

Sociodemographic (self-report), cognitive, and psychiatric data was assessed through the COllaborative Informatics and Neuroimaging Suite (COINS) Data Exchange after completion of appropriate data use agreements. We selected a number of measures that we believed may covary with T1-weighted MRI quality. Motivated by past studies, these included: age, sex, body mass index (BMI), general cognitive ability (IQ), and clinical diagnoses. The Wechsler Intelligence Scale for Children (WISC-V) was used as a measure of general cognitive ability (IQ) and was completed on 336 participants in the sample; the WISC-V is an individually administered clinical instrument for assessing the intelligence of youth participants 6-16 and generates a general cognitive ability score (Full-Scale Intelligence Quotient; FSIQ). Related to clinical diagnoses, the presence of psychopathology was assessed by a certified clinician using semi-structured DSM-5-based psychiatric interview (i.e., the Schedule for Affective Disorders and Schizophrenia for Children; KSADS-COMP). This data was available for 367 participants in our sample. Mean, standard deviation, and ranges for all the sociodemographic, cognitive, and psychiatric measures are noted in Table 1. Additional information about these measures is noted in our *Supplemental Materials*.

### 3.1 Image Pre/processing (Freesurfer)

Standard-processing approaches from Freesurfer (e.g., cortical reconstruction; volumetric segmentation) were performed in version 7.1. Freesurfer is a widely-documented and freely available morphometric processing tool suite (http://surfer.nmr.mgh.harvard.edu/). The technical details of these procedures are described in prior publications [26–31]. Briefly, this processing includes motion correction and intensity normalization of T1-weighted images, removal of non-brain tissue using a hybrid watershed/surface deformation procedure [32], automated Talairach transformation, segmentation of the subcortical white matter and deep gray matter volumetric structures (including hippocampus, amygdala, caudate, putamen, ventricles), tessellation of the gray matter white matter boundary, and derivation of cortical surface area and cortical thickness. Of note, the “recon-all” pipeline with the default set of parameters (no flag options) was used and no manual editing was conducted. After successful processing, we extracted volumes from subcortical structures, as well as mean cortical surface area and cortical thickness for the 34 bilateral Desikan Killiany (DK) atlas regions [33]. Freesurfer was implemented using Brainlife.io, (brainlife.app.0, https://doi.org/10.25663/bl.app.0), which is a free, publicly funded, cloud-computing platform for reproducible neuroimaging pipelines and data sharing [34] also, for additional information, visit http://brainlife.io/). Scans from four participants did not complete processing in Freesurfer due to technical issues; this brought the total sample size that passed visual inspection and with Freesurfer processing completed) to N=205. Graphically depictions of our methods are shown in Figure 1.

**Figure 1.**
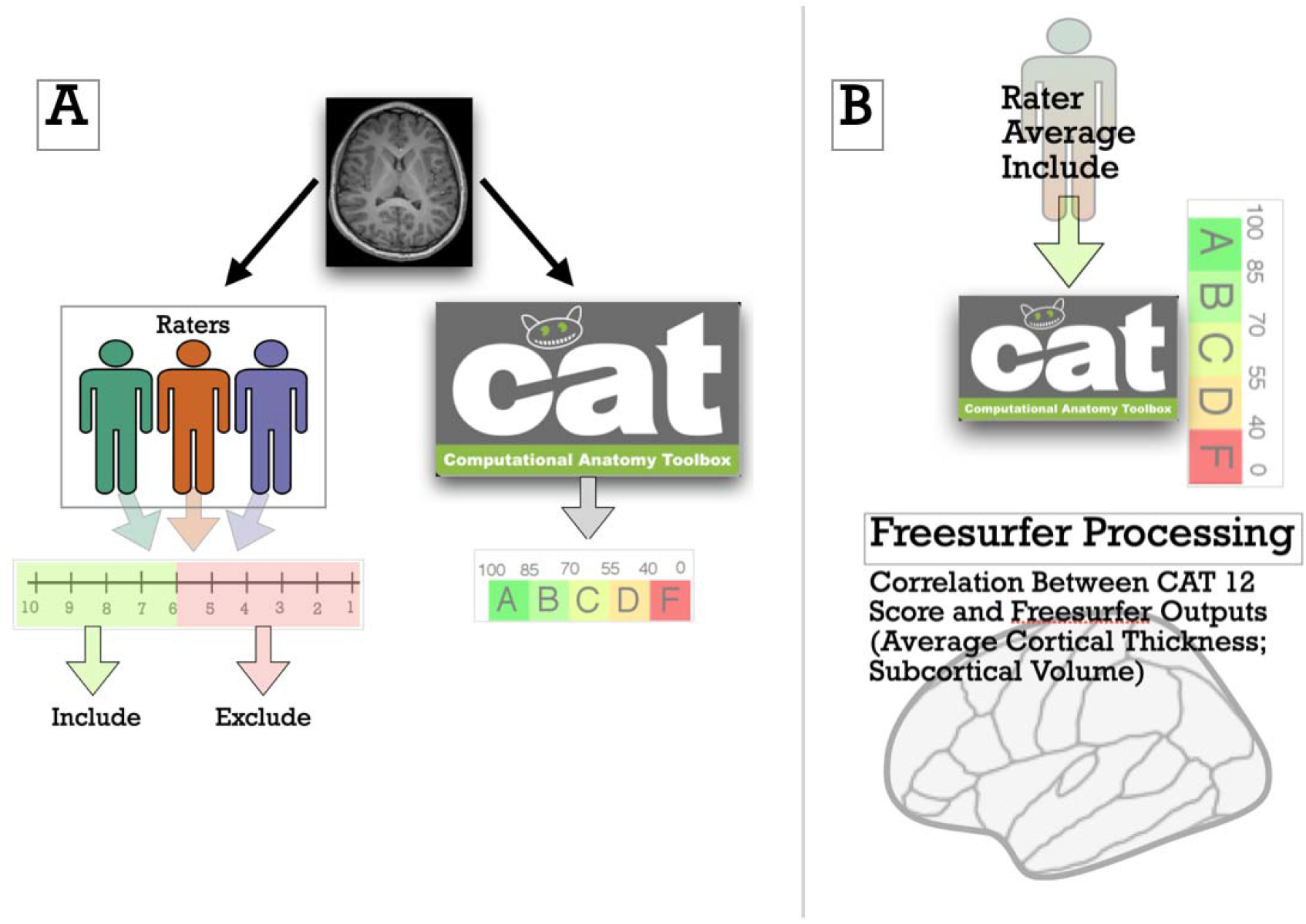
Graphic depiction of the study’s procedures. Structural MRI images were rated by multiple trained research assistant and also processed in the CAT12 toolbox (Panel A). Human Raters rated each image and then these ratings were averaged; MRI images, with a rating <u>>6, were then processed in Freesurfer, and relations between CAT12 scores and Freesurfer outputs were examined (Panel B)</u>.

### 3.2 Statistical Modeling

We first constructed logistic regression models that used an aggregated measure of T1-weighted image quality from the CAT12 toolbox and the outcome of passing or failing visual quality assurance checks completed by trained human raters. Receiver operating characteristic curves were computed to understand true positive (sensitivity) and false positive rates. For these receiver operating characteristic measures, the area under the curve (AUC) was computed to show classification performance at all classification thresholds (and distinguishing between classes of passing or failing visual quality assurance checks). We additionally constructed: 1) Bayesian logistic models, and 2) Confusion matrices. Bayesian logistic models probed potential over-fitting and biases common to Frequentist logistic models [35]. Confusion matrix construction involved logistic model fitting on 80% of our full sample (as a “training” set) and then application of these parameters to the remaining 20% of our sample (the “test” set). Next, bivariate correlations were calculated to examine relations between our image quality and sociodemographic variables of interest, including age, sex, IQ, BMI, and clinical diagnosis. Finally, we computed 158 bivariate correlations between T1-weighted image quality and Freesurfer outputs (68 mean cortical surface area from the DK atlas; 68 mean cortical thickness estimates from the DK atlas; 22 subcortical regions). Of note, cerebral spinal fluid Freesurfer subcortical outputs (e.g., Lateral ventricle; Left-choroid-plexus) were excluded from analyses.

Given the number of statistical tests conducted and to further reproducibility, we adjusted all p-values of this last step based on the Benjamini & Hochberg False Discovery Rate Correction [36]. This commonly-used approach has been shown to have appropriate power to detect true positives, while still controlling the proportion of type I errors at a specified level (α=.05). This was done “within” each morphometric output categories (i.e., correcting for 68 correlations for surface area and MRI quality). We graphed all results with ‘*ggseg’* R library [37]. All reported correlations are derived from linear regression models with 1 independent variable, so this can be seen as equivalent to a bivariate (Pearson’s) correlation coefficient. A pdf version of our RMarkdown output is available in our Supplemental Materials and online (at https://github.com/jlhanson5/Gilmore_Buser_Hanson_CAT12_Freesurfer).

### 3.3 Supplemental Modeling

To probe the robustness of the results reported in the main document, we also completed a number of follow-up analyses related to our variables of interest. These included: 1) Constructing logistic regression models and ROC curves with CAT12 and another marker of image quality, Freesurfer’s Euler number; 2) Examining associations between Freesurfer outputs and structural MRI quality after controlling for the important sociodemographic factor of age; 3) Testing associations between Freesurfer outputs and CAT12 scores, after controlling for Freesurfer’s Euler number; and 4) Probing relations between Freesurfer outputs and CAT12 scores in participants excluded after visual quality checks. Please see our *Supplemental Materials* for these additional analyses.

## 4. RESULTS

### 4.1 Relations Between T1-weighted MRI Quality and Visual Rejection/Acceptance of Structural Images

Logistic regression was used to examine relationships between our aggregated T1-weighted MRI quality measure and the outcome of passing or failing quality assurance checks completed by trained human raters. Logistic regression models indicated that T1-weighted MRI quality, derived by the CAT12 toolbox, was significantly related to passing or failing quality assurance checks completed by trained human raters (*z=7*.*877, p<*.*005; Nagelkerke’s R*^*2*^*=0*.*8951)*. This indicated that greater CAT12 MRI quality scores were related to a higher likelihood of passing visual inspection. Receiver operating characteristic analyses indicated a mean AUC of 98.9% (with 95% confidence intervals spanning 98.2-99.6%, as shown in Figure 2). Bayesian GLM modeling suggested a similar relation, with higher MRI quality significantly relating to passing visual checks (*z=8*.*141, p<*.*005)*. As shown in Figure 3, Confusion matrices indicated strong model prediction, out of sample (derived from 80% of our sample, to a heldout 20%)--Accuracy=0.937 and Kappa=0.869.

**Figure 2.**
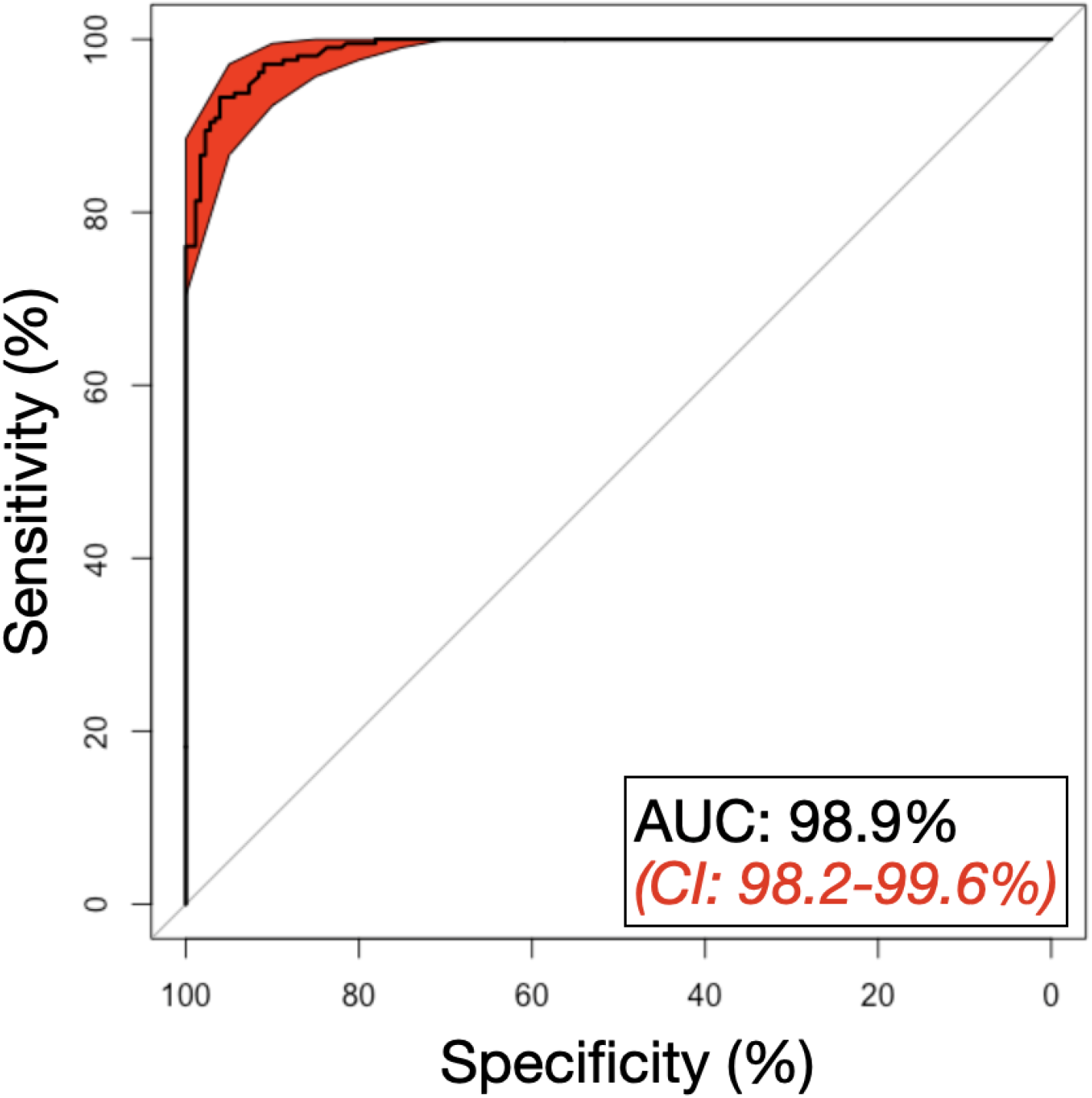
ROC curves showing the validity of image quality (derived from the CAT12 toolbox) for discriminating passing (versus failing) human rater visual checks of quality. Sensitivity and specificity were both high, suggesting image quality was able to robustly parse this binary categorization. 95% Confidence Intervals of these ROC curves are shown in red.

**Figure 3.**
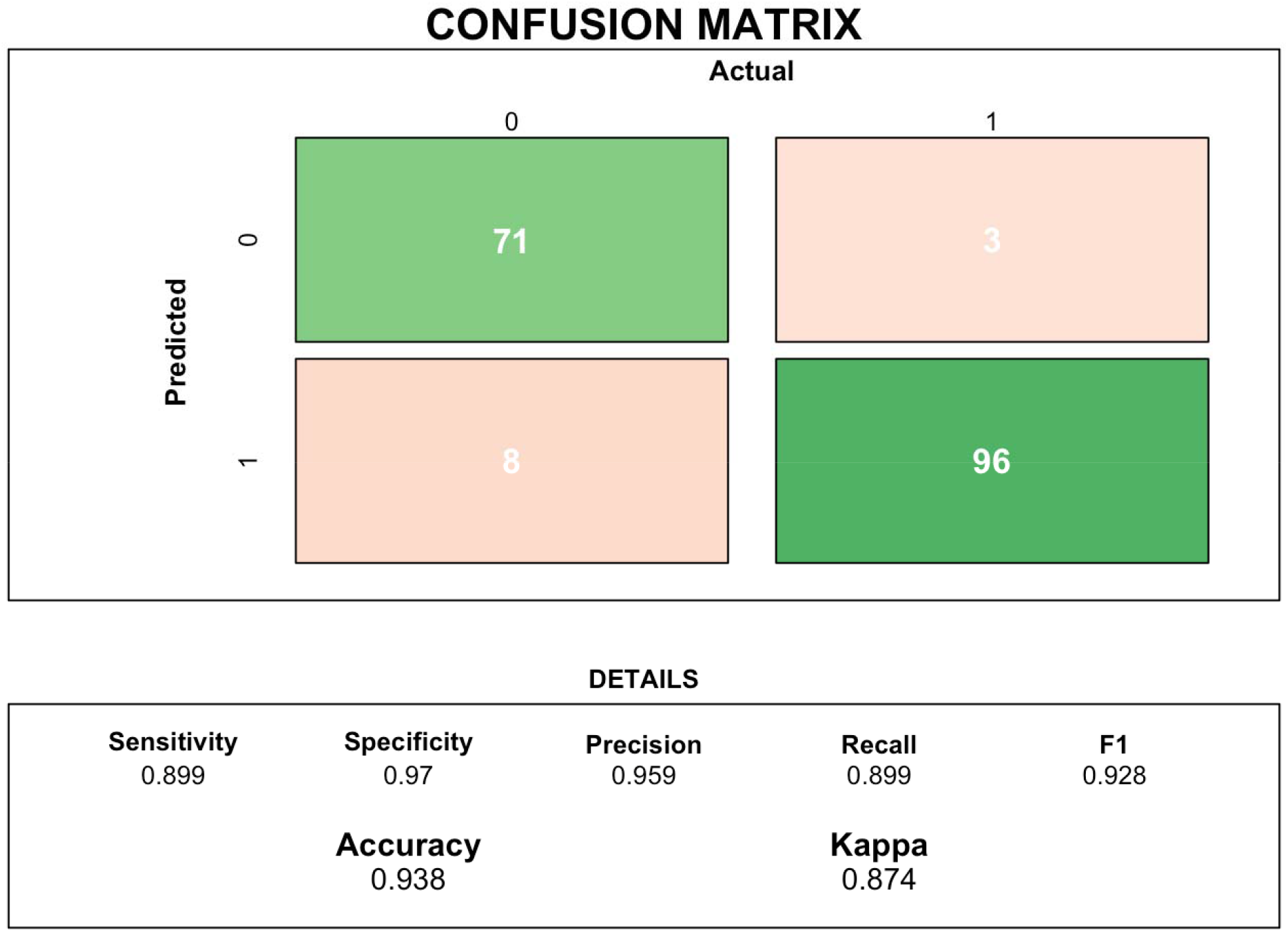
To further probe the ability of CAT12 scores to accurately classified inclusion/exclusion of MRI images (derived from our human raters), confusion matrices were constructed. Of note, 80% of our data was used in our training set and 20% in our test set. This graphic displays the different metric of accurate classification including sensitivity, specificity, accuracy, and kappa.

### 4.2 Bivariate Correlations Between T1-weighted Image Quality and Sociodemographic Variables of Interest

We next examined correlations between T1-weighted image quality, sociodemographic variables of interest (e.g., age, sex, BMI, and clinical diagnosis). As expected and in line with other reports, image quality was related to age (r=0.321, p<.005; as shown in Figure 2). Older subjects typically had better quality scans. Interestingly, no other sociodemographic factors were significantly related to image quality (Sex p=0.196; BMI p=.227; Clinical Diagnosis [binary indicator] p=.189). The BMI finding is in contrast to past results reported in adults [8, 38]. There was a trend association for image quality and IQ (r=.101, p= 0.06), with high IQ relating to better image quality. Of note, this is for all participants (not only those passing human rater visual inspection). If associations are investigated in only those passing visual inspection, the association with age and image quality remains significant (p=0.036). All other associations remained non-significant (all p’s >.3).

**Figure 4.**
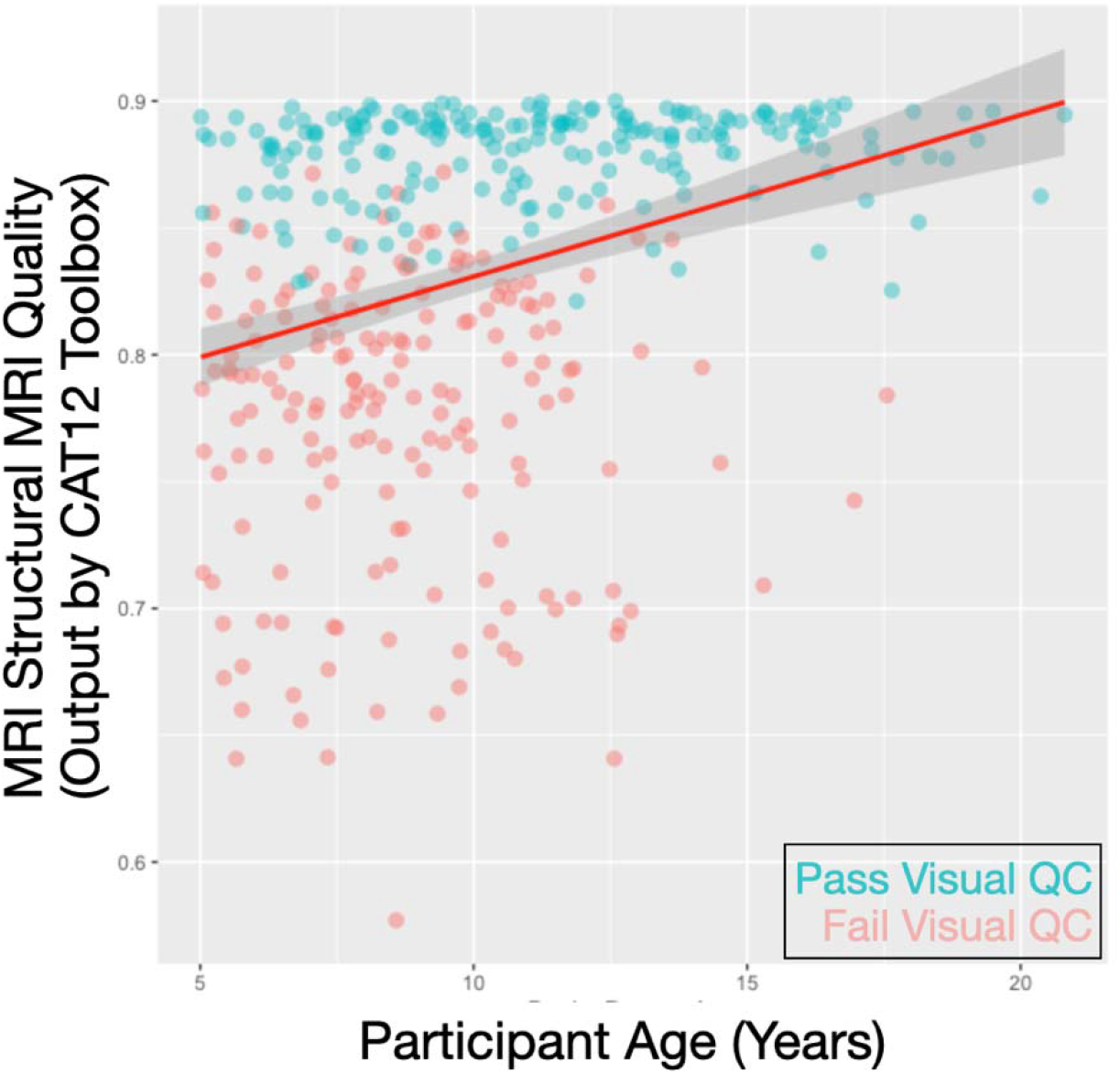
Scatterplot showing participant age (in years; horizontal axis) and image quality (an aggregated measure of noise to contrast ratio, coefficient of joint variation, inhomogeneity to contrast ratio, and Root mean squared voxel resolution, ranging from 0-1; vertical axis). Dot color indicates whether the participants passed visual quality checks (Pass = Turquoise ; Fail = Salmon)

### 4.3 Associations Between Freesurfer Outputs and Structural MRI Quality

We next examined correlations between T1-weighted MRI quality and 158 morphometric outputs from Freesurfer (68 mean cortical surface area estimates from the DK atlas; 68 mean cortical thickness estimates also from the DK atlas; 22 subcortical regions). Related to cortical surface area, there was variability in how T1-weighted image quality related to mean surface area from differ brain parcels (t-statistic range=-0.926-4.918). In aggregate, this association was modest (Mean t-statistic=1.473 +/- 1.33); however, in twelve areas, the association between image quality and mean surface area was significant, even after correcting for multiple comparisons (p _fdr-corrected_<.05, as displayed in Table 2 and Figure 5). For cortical thickness, there was again variability in relation between mean thickness for parcels and image quality (t-statistic range=-2.376-6.571), with modest associations in the aggregate (Mean t-statistic=1.510 +/- 2.04). However, relations between image quality and cortical thickness for twenty-three regions was significant, even after correcting for multiple comparisons (p _fdr-corrected_<.05, as shown in Table 3 and Figure 6). Finally, for subcortical volume, similar patterns were seen (t-statistic range=-0.5896-3.337; mean t-statistic=1.312 +/- 1.016, as shown in Table 3 and Figure 6). Of note, volumes from two regions, the left amygdala and the posterior portion of the corpus callosum, were related to image quality (p _fdr-corrected_<.05) after correcting for multiple comparisons. In the aggregate, we examined 158 morphometric outputs from Freesurfer and 37 were significantly related to image quality, after correcting for multiple comparisons. Of note, if one did not correct for multiple comparisons, 56 regions (or ∼35.4% of the outputs) were related to image quality at p<.05.

**Table 2.** Relations Between Cortical Surface Area and Structural MRI Quality (as measured by the CAT12 Toolbox). Table displays relations between MRI quality (CAT12 score) and cortical surface area for different brain parcels in Freesurfer’s DK atlas. The left side of the table shows regions in the left hemisphere, while the right side shows the right hemisphere. On each side, region is in the first column, and t-statistic (of CAT12 and cortical surface area) is in the second column. The third column is the uncorrected p-value, while the fourth column is this test statistic corrected for multiple comparisons (for all 68 cortical parcels). Light orange highlighting indicates regions that were p<.05 (*uncorrected*), while darker orange highlighting indicates regions that were p<.05 (*FDR corrected*).

| Area Parcel | t_statistics | p_value | p_adjusted |
| --- | --- | --- | --- |
| lh_bankssts | 0.44371 | 0.65772 | 0.79866 |
| lh_caudalanteriorcingulate | 2.08777 | 0.03805 | 0.11760 |
| lh_caudalmiddlefrontal | 1.17122 | 0.24286 | 0.48195 |
| lh_cuneus | -0.44706 | 0.65530 | 0.79866 |
| lh_entorhinal | 1.22860 | 0.22062 | 0.45462 |
| lh_fusiform | 0.81376 | 0.41672 | 0.59110 |
| lh_inferiorparietal | 0.30567 | 0.76016 | 0.86025 |
| lh_inferiortemporal | 4.10177 | 0.00006 | 0.00080 |
| lh_isthmuscingulate | 0.18245 | 0.85541 | 0.89489 |
| lh_lateraloccipital | -0.61571 | 0.53877 | 0.73272 |
| lh_lateralorbitofrontal | 4.36176 | 0.00002 | 0.00046 |
| lh_lingual | 0.13832 | 0.89012 | 0.91709 |
| lh_medialorbitofrontal | 3.23941 | 0.00140 | 0.01357 |
| lh_middletemporal | 2.61381 | 0.00961 | 0.05029 |
| lh_parahippocampal | 1.09311 | 0.27562 | 0.49322 |
| lh_paracentral | 0.73901 | 0.46074 | 0.63940 |
| lh_parsopercularis | 0.81284 | 0.41725 | 0.59110 |
| lh_parsorbitalis | 2.54329 | 0.01172 | 0.05311 |
| lh_parstriangularis | 2.14037 | 0.03350 | 0.11390 |
| lh_pericalcarine | 0.23182 | 0.81691 | 0.88174 |
| lh_postcentral | -0.35881 | 0.72011 | 0.85907 |
| lh_posteriorcingulate | 2.88885 | 0.00428 | 0.02910 |
| lh_precentral | -0.92631 | 0.35537 | 0.56197 |
| lh_precuneus | 2.04056 | 0.04257 | 0.12586 |
| lh_rostralanteriorcingulate | 3.08607 | 0.00231 | 0.01961 |
| lh_rostralmiddlefrontal | 4.67811 | 0.00001 | 0.00018 |
| lh_superiorfrontal | 1.95419 | 0.05203 | 0.13748 |
| lh_superiorparietal | 1.06070 | 0.29007 | 0.49980 |
| lh_superiortemporal | 0.94196 | 0.34732 | 0.56197 |
| lh_supramarginal | 1.46012 | 0.14578 | 0.34183 |
| lh_frontalpole | 2.33001 | 0.02077 | 0.07848 |
| lh_temporalpole | 2.11599 | 0.03555 | 0.11510 |
| lh_transversetemporal | 0.85295 | 0.39468 | 0.59110 |
| lh_insula | 2.37299 | 0.01856 | 0.07426 |
| rh_bankssts | 0.83044 | 0.40725 | 0.59110 |
| rh_caudalanteriorcingulate | 0.85123 | 0.39563 | 0.59110 |
| rh_caudalmiddlefrontal | 1.93317 | 0.05459 | 0.13748 |
| rh_cuneus | -0.18496 | 0.85344 | 0.89489 |
| rh_entorhinal | 1.25415 | 0.21121 | 0.44882 |
| rh_fusiform | 1.93642 | 0.05418 | 0.13748 |
| rh_inferiorparietal | 0.46049 | 0.64565 | 0.79866 |
| rh_inferiortemporal | 3.35107 | 0.00096 | 0.01085 |
| rh_isthmuscingulate | 1.05208 | 0.29400 | 0.49980 |
| rh_lateraloccipital | 0.55048 | 0.58259 | 0.76184 |
| rh_lateralorbitofrontal | 4.10849 | 0.00006 | 0.00080 |
| rh_lingual | -0.25066 | 0.80233 | 0.87997 |
| rh_medialorbitofrontal | 4.91858 | 0.00000 | 0.00012 |
| rh_middletemporal | 2.29156 | 0.02294 | 0.08211 |
| rh_parahippocampal | 0.29054 | 0.77169 | 0.86025 |
| rh_paracentral | 0.96544 | 0.33546 | 0.55636 |
| rh_parsopercularis | 1.15082 | 0.25114 | 0.48195 |
| rh_parsorbitalis | 2.56615 | 0.01099 | 0.05311 |
| rh_parstriangularis | 0.04814 | 0.96165 | 0.96165 |
| rh_pericalcarine | -0.09957 | 0.92079 | 0.93453 |
| rh_postcentral | 1.38743 | 0.16681 | 0.37810 |
| rh_posteriorcingulate | 1.30056 | 0.19486 | 0.42744 |
| rh_precentral | 0.33314 | 0.73937 | 0.86025 |
| rh_precuneus | 2.43302 | 0.01583 | 0.06726 |
| rh_rostralanteriorcingulate | 2.91711 | 0.00392 | 0.02910 |
| rh_rostralmiddlefrontal | 2.67074 | 0.00817 | 0.04832 |
| rh_superiorfrontal | 1.90633 | 0.05800 | 0.14085 |
| rh_superiorparietal | 0.55646 | 0.57850 | 0.76184 |
| rh_superiortemporal | 1.09531 | 0.27466 | 0.49322 |
| rh_supramarginal | 2.65600 | 0.00853 | 0.04832 |
| rh_frontalpole | 0.47238 | 0.63715 | 0.79866 |
| rh_temporalpole | 1.14111 | 0.25515 | 0.48195 |
| rh_transversetemporal | -0.32295 | 0.74706 | 0.86025 |
| rh_insula | 1.96440 | 0.05083 | 0.13748 |

**Table 3.** Relations Between Cortical Thickness and Structural MRI Quality (as measured by the CAT12 Toolbox). Table displays relations between MRI quality (CAT12 score) and cortical thickness for different brain parcels in Freesurfer’s DK atlas. The left side of the table shows regions in the left hemisphere, while the right side shows the right hemisphere. On each side, region is in the first column, and t-statistic (of CAT12 and cortical thickness) is in the second column. The third column is the uncorrected p-value, while the fourth column is this test statistic corrected for multiple comparisons (for all 68 parcels). Light orange highlighting indicates regions that were p<.05 (*uncorrected*), while darker orange highlighting indicates regions that were p<.05 (*FDR corrected*).

| Thickness Parcel | t_statistics | p_value | p_adjusted | Thickness Parcel | t_statistics | p_value | p_adjusted |
| --- | --- | --- | --- | --- | --- | --- | --- |
| lh_bankssts | 1.08997 | 0.27700 | 0.40948 | rh_bankssts | 1.55943 | 0.12043 | 0.20998 |
| lh_caudalanteriorcingulate | -1.34432 | 0.18032 | 0.29907 | rh_caudalanteriorcingulate | -0.57201 | 0.56794 | 0.66586 |
| lh_caudalmiddlefrontal | 4.30970 | 0.00003 | 0.00022 | rh_caudalmiddlefrontal | 4.45687 | 0.00001 | 0.00013 |
| lh_cuneus | 0.15952 | 0.87342 | 0.91373 | rh_cuneus | 0.89066 | 0.37415 | 0.50884 |
| lh_entorhinal | 5.07313 | 0.00000 | 0.00001 | rh_entorhinal | 5.28642 | 0.00000 | 0.00001 |
| lh_fusiform | 2.68890 | 0.00776 | 0.02526 | rh_fusiform | 2.58405 | 0.01046 | 0.03091 |
| lh_inferiorparietal | 2.74840 | 0.00652 | 0.02463 | rh_inferiorparietal | 1.28643 | 0.19974 | 0.30869 |
| lh_inferiortemporal | 2.60408 | 0.00988 | 0.03055 | rh_inferiortemporal | 2.89207 | 0.00424 | 0.02058 |
| lh_isthmuscingulate | -1.90339 | 0.05838 | 0.12807 | rh_isthmuscingulate | -2.20428 | 0.02861 | 0.07483 |
| lh_lateraloccipital | 2.86879 | 0.00455 | 0.02062 | rh_lateraloccipital | 2.78964 | 0.00577 | 0.02372 |
| lh_lateralorbitofrontal | 2.02652 | 0.04400 | 0.10011 | rh_lateralorbitofrontal | 2.17463 | 0.03080 | 0.07756 |
| lh_lingual | -1.65309 | 0.09983 | 0.18348 | rh_lingual | -0.63065 | 0.52897 | 0.65336 |
| lh_medialorbitofrontal | -0.23206 | 0.81673 | 0.88155 | rh_medialorbitofrontal | -2.37653 | 0.01839 | 0.05211 |
| lh_middletemporal | 2.26172 | 0.02476 | 0.06734 | rh_middletemporal | 2.78051 | 0.00593 | 0.02372 |
| lh_parahippocampal | 0.60225 | 0.54767 | 0.65336 | rh_parahippocampal | 2.02490 | 0.04417 | 0.10011 |
| lh_paracentral | 1.85395 | 0.06518 | 0.13850 | rh_paracentral | 2.68693 | 0.00780 | 0.02526 |
| lh_parsopercularis | 1.76791 | 0.07856 | 0.15382 | rh_parsopercularis | 2.71333 | 0.00722 | 0.02526 |
| lh_parsorbitalis | 1.22890 | 0.22051 | 0.33322 | rh_parsorbitalis | 0.12045 | 0.90424 | 0.91774 |
| lh_parstriangularis | 1.00086 | 0.31807 | 0.45060 | rh_parstriangularis | 0.75355 | 0.45198 | 0.59105 |
| lh_pericalcarine | -1.57667 | 0.11641 | 0.20831 | rh_pericalcarine | -0.14035 | 0.88852 | 0.91545 |
| lh_postcentral | 1.76425 | 0.07917 | 0.15382 | rh_postcentral | 0.64465 | 0.51987 | 0.65336 |
| lh_posteriorcingulate | -0.24391 | 0.80755 | 0.88155 | rh_posteriorcingulate | -1.05990 | 0.29043 | 0.42020 |
| lh_precentral | 6.57110 | 0.00000 | 0.00000 | rh_precentral | 5.57632 | 0.00000 | 0.00000 |
| lh_precuneus | 0.61377 | 0.54005 | 0.65336 | rh_precuneus | 0.38736 | 0.69889 | 0.79208 |
| lh_rostralanteriorcingulate | -1.52622 | 0.12849 | 0.21843 | rh_rostralanteriorcingulate | 0.49376 | 0.62200 | 0.71688 |
| lh_rostralmiddlefrontal | -0.18503 | 0.85339 | 0.90672 | rh_rostralmiddlefrontal | -0.09858 | 0.92157 | 0.92157 |
| lh_superiorfrontal | 1.77457 | 0.07745 | 0.15382 | rh_superiorfrontal | 1.29407 | 0.19709 | 0.30869 |
| lh_superiorparietal | 2.90099 | 0.00412 | 0.02058 | rh_superiorparietal | 2.97047 | 0.00333 | 0.02056 |
| lh_superiortemporal | 4.18747 | 0.00004 | 0.00032 | rh_superiortemporal | 2.89752 | 0.00417 | 0.02058 |
| lh_supramarginal | 3.78244 | 0.00020 | 0.00138 | rh_supramarginal | 1.71206 | 0.08839 | 0.16696 |
| lh_frontalpole | -1.29074 | 0.19824 | 0.30869 | rh_frontalpole | -0.28271 | 0.77768 | 0.86692 |
| lh_temporalpole | 5.46017 | 0.00000 | 0.00000 | rh_temporalpole | 5.11553 | 0.00000 | 0.00001 |
| lh_transversetemporal | 0.81222 | 0.41760 | 0.55680 | rh_transversetemporal | 2.07574 | 0.03916 | 0.09510 |
| lh_insula | 0.97027 | 0.33305 | 0.46219 | rh_insula | 0.71959 | 0.47259 | 0.60635 |

**Table 4.** Relations Between Subcortical Volumes and Structural MRI Quality (as measured by the CAT12 Toolbox). Table displays relations between MRI quality (CAT12 score) and subcortical volumes in Freesurfer’s ASEG atlas. Region is in the first column, and t-statistic (of CAT12 and subcortical volume) is in the second column. The third column is the uncorrected p-value, while the fourth column is this test statistic corrected for multiple comparisons (for all 22 subcortical regions of interest). Light orange highlighting indicates regions that were p<.05 (*uncorrected*), while darker orange highlighting indicates regions that were p<.05 (*FDR corrected*).

| Aseg_Volume | t_statistics | p_value | p_adjusted |
| --- | --- | --- | --- |
| Left-Thalamus-Proper | -0.10352 | 0.91765 | 0.97864 |
| Left-Caudate | 0.18568 | 0.85288 | 0.97864 |
| Left-Putamen | 1.64437 | 0.10163 | 0.22358 |
| Left-Pallidum | 1.32804 | 0.18563 | 0.31415 |
| brain-stem | 1.39762 | 0.16373 | 0.30017 |
| Left-Hippocampus | 0.87275 | 0.38381 | 0.55460 |
| Left-Amygdala | 3.26946 | 0.00126 | 0.01389 |
| Left-Accumbens-area | 0.02680 | 0.97864 | 0.97864 |
| Left-VentralDC | 0.82099 | 0.41260 | 0.55460 |
| Right-Thalamus-Proper | -0.58969 | 0.55604 | 0.67961 |
| Right-Caudate | 0.07730 | 0.93846 | 0.97864 |
| Right-Putamen | 1.71325 | 0.08817 | 0.21891 |
| Right-Pallidum | 1.74292 | 0.08284 | 0.21891 |
| Right-Hippocampus | 0.79323 | 0.42856 | 0.55460 |
| Right-Amygdala | 1.89779 | 0.05912 | 0.21678 |
| Right-Accumbens-area | 1.48388 | 0.13937 | 0.27874 |
| Right-VentralDC | 1.01197 | 0.31274 | 0.49144 |
| cc-posterior | 3.33701 | 0.00100 | 0.01389 |
| cc-mid-posterior | 1.70579 | 0.08956 | 0.21891 |
| cc-central | 1.96198 | 0.05111 | 0.21678 |
| cc-mid-anterior | 2.33941 | 0.02027 | 0.14866 |
| cc-anterior | 1.96509 | 0.05075 | 0.21678 |

**Figure 5.**
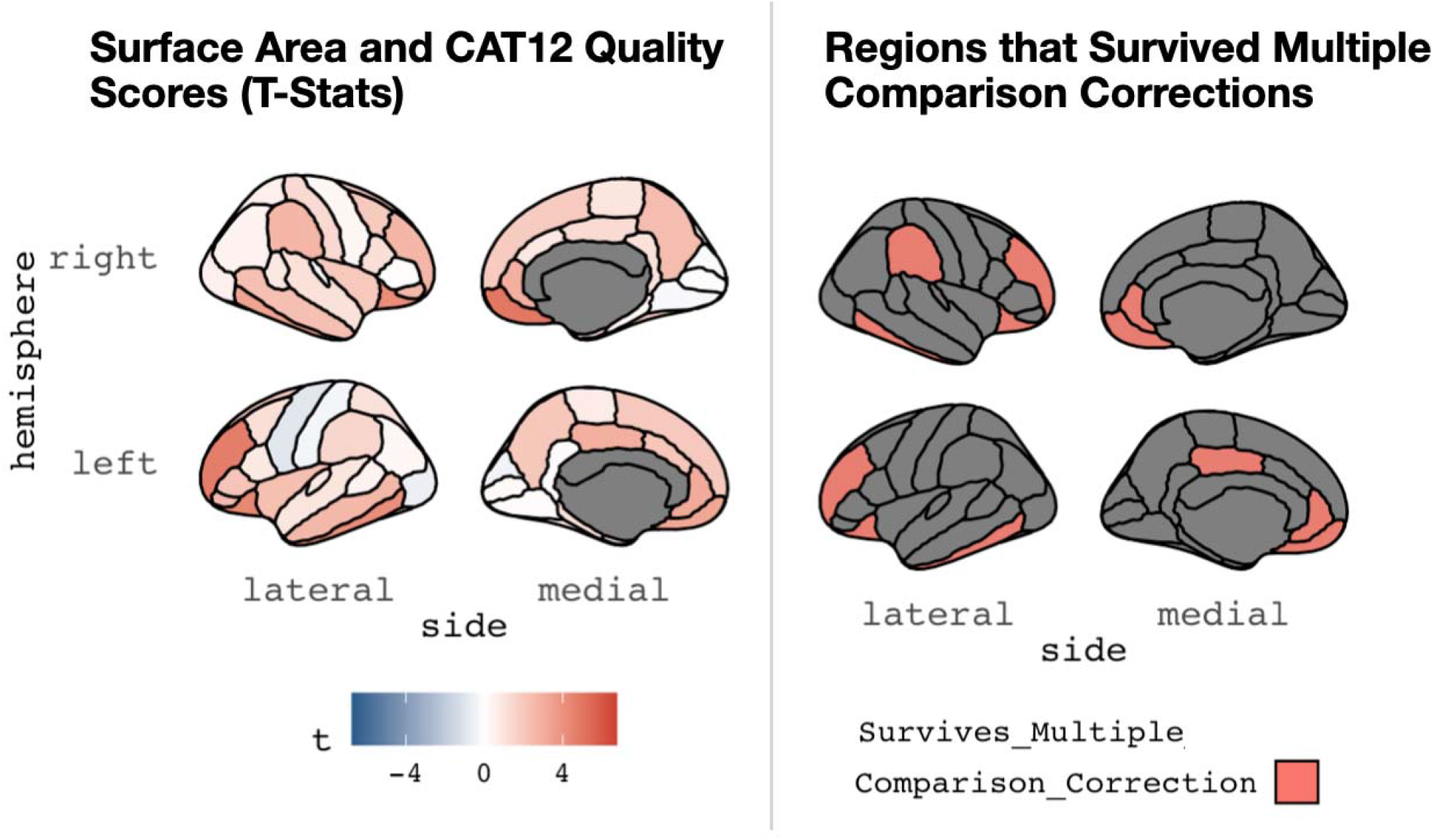
A graphic depiction (from the R library *ggseg*) showing associations between image quality (assessed by the CAT12 Toolbox) and derived (mean) cortical surface area. This is shown for the DK atlas commonly used in Freesurfer. Lateral and medial views are shown for the right (top) and left (bottom) hemispheres. The left panel shows the overall t-statistics for the relation in each parcel, while the right panel shows parcels where the relation between surface area and image quality survives multiple comparisons.

**Figure 6.**
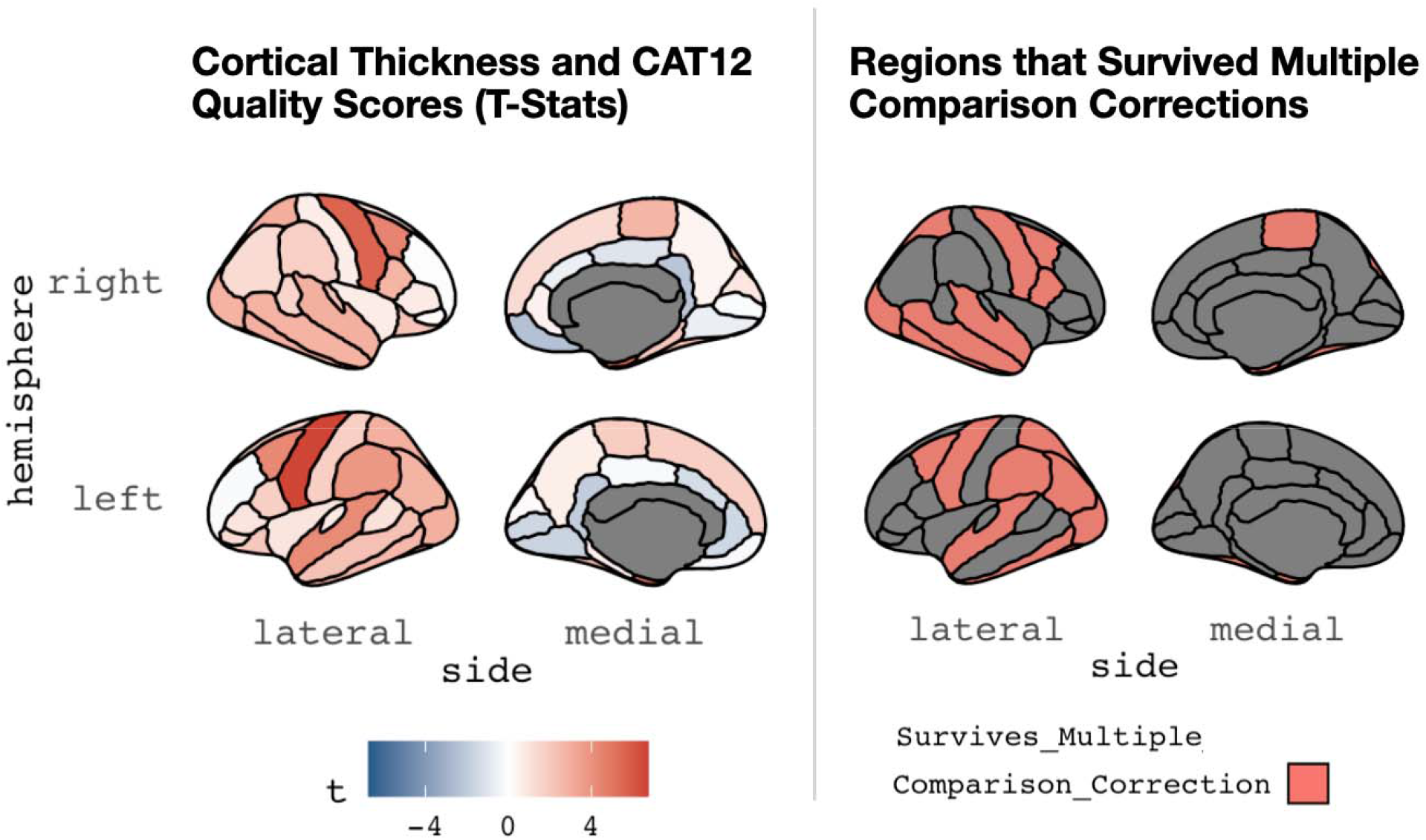
A graphic depiction (from the R library *ggseg*) showing associations between image quality (assessed by the CAT12 Toolbox) and derived (mean) cortical thickness. This is shown for the DK atlas commonly used in Freesurfer. Lateral and medial views are shown for the right (top) and left (bottom) hemispheres. The left panel shows the overall t-statistics for the relation in each parcel, while the right panel shows parcels where the relation between cortical thickness and image quality survives multiple comparisons.

**Figure 7.**
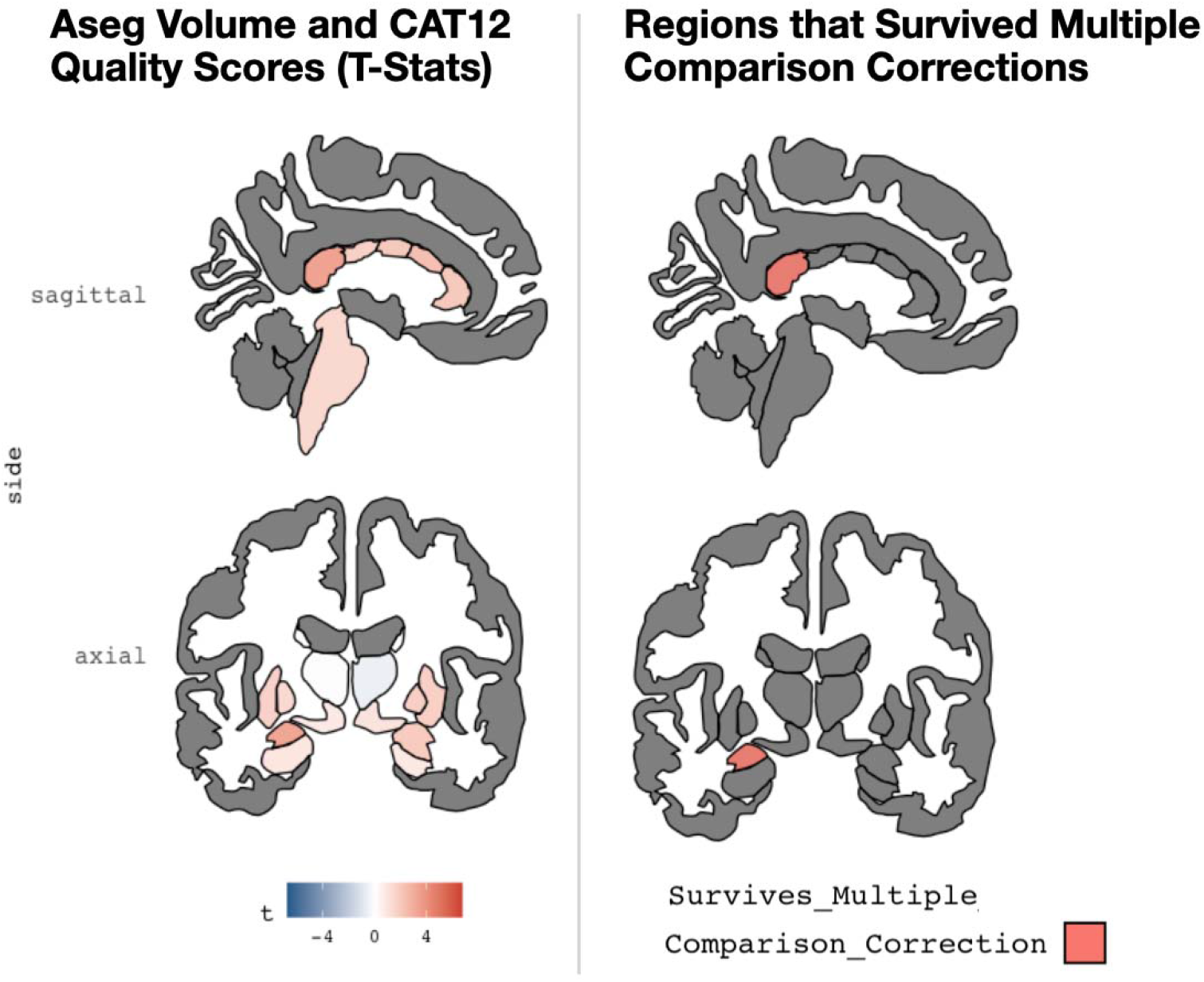
A graphic depiction (from the R library ggseg) showing associations between image quality (assessed by the CAT12 Toolbox) and subcortical volumes. This is shown for the Freesurfer ASEG atlas. Coronal (left) and sagittal (right) views are shown. The left panel shows the t-statistic for the relation in each subcortical volume, while the right panel shows parcels where the relation between volume and image quality survives multiple comparisons.

## 5. DISCUSSION

The primary goals of this study were three-fold: 1) to see if an integrated measure of image quality (output by the CAT12 toolbox) related to visual rater judgement (retain/exclude) of T1-weighted MRI images; 2) to examine if direct measures of T1-weighted imaging quality were associated with sociodemographic and behavioral variables of interest; 3) to investigate if there were associations between commonly-used Freesurfer outputs and T1-weighted image quality. Related to the first goal (and perhaps as expected), the measure of image quality output by the CAT12 toolbox was strongly related to visual rater judgement of T1-weighted MRI images. Logistic regression models and receiver operating characteristic analyses supported this idea. Connected to this second goal, we found significant associations between image quality and age; there were, however, no relations between IQ, BMI, sex, or clinical diagnosis. Finally, we demonstrated commonly-derived structural MRI measures, derived from T1-weighted images, were strongly related to image quality. Even after correcting for multiple comparisons, numerous measurements of cortical surface area, cortical thickness and subcortical volumes were connected to image quality. This was for a large percentage (23.4%) of the brain regions investigated, suggesting diffuse, but significant, impacts of image quality on structural morphometric measures. Interestingly, many of the regions that survive multiple comparisons (e.g., entorhinal, precentral, caudal middle frontal parcels) were found to be influential in the automated quality control suite, Qoala-T [16]. Examined collectively, our results have significant implications for studies of neurodevelopment and other applied work using T1-weighted MRI, as motion artifacts are especially problematic for young children and clinical populations; these groups may have difficulty remaining still during the time required to collect high-resolution neuroimaging data.

Contextualizing our results with past research reports, we find significant bivariate associations between image quality and age. However, we did not find associations between image quality and factors such as general intelligence (IQ), and BMI. Such findings are in contrast to a few prior publications [8, 38]. This may be due to the age-range of our sample (5-21 years of age), while those relevant past studies have been primarily completed in adult samples. Building off of previous studies, we find image quality is related to derived measures of brain anatomy, irrespective of typical (binary) quality threshold cut-offs. Even in structural scans of high quality (that “pass” visual inspection), in-scanner motion appears to influence morphometric estimations. Indeed, accurate quantification of regional grey matter volume relies on reliable segmentation from high-resolution MR images. Head motion during an MRI scan can bias segmentation, which in turn can impact morphometric measurements.

Our results have important implications when thinking about structural MRI, especially for studies attempting to center-in on individual differences using T1-weighted MR scans. We used more direct measures of T1-weighted image quality, rather than measures derived from resting state (e.g., Refs.[14, 17]). Use of resting state may capture aspects of participant movement, but it is not specifically during the T1-weighted MRI scan. Furthermore, this type of information may not be available for all studies, but the measure we employ here could be derived for any T1-weighted scan. Using this more direct measure of MRI quality, we found impacts on morphometric variables typically generated from T1-weighted MR images. For example, other studies have used proxy measures for image quality derived from subject-motion during functional scans [12, 17]. However, proxy measures for subject motion may be missing true differences obscured by motion [20]. Our findings build off of past work by Rosen and colleagues’ that found Freesurfer Euler number was related to Freesurfer cortical thickness measures. Here, however, we used a more direct metric of image quality, derived from the CAT12 toolbox, and examined correlations with this measure and commonly-used Freesurfer outputs. This use of an independent image quality metric provides stronger evidence of the impact of image quality on subcortical volume, cortical surface area, and cortical thickness. Across these Freesurfer outputs, there was variability in image quality and relations with surface area, thickness and volumes; positive and negative relations were commonly noted across the different atlases. However, the only relations between image quality and Freesurfer outputs that survived multiple comparisons were positive in nature--greater image quality related to higher values in these regions. Interestingly, many of the regions that survive multiple comparisons (e.g., entorhinal, precentral, caudal middle frontal parcels) were found to be influential in the automated quality control suite, Qoala-T [16]. These areas may be particularly impacted by participant motion and image quality. Finally, and of interest to those studying emotion, we find that volumetric measures of amygdala were related to image quality, with higher image quality relating to higher volumes in this area.

Considering our project, as well as past studies, our results suggest it will be important to consider image quality in future structural MRI analyses using T1-weighted images. In line with current work, studies interested in individual and/or group differences should flag/exclude scans of extremely poor quality. Furthermore, in the future, research groups may think about accounting for individual differences in motion-related image quality by using more direct measures of image quality as covariates in morphometric analyses. Such a strategy could address indirect effects of motion-related image quality and to confirm main effects for their variables of interest. However, as with any covariate of “*no interest*”, if motion is collinear with other variables, important variance related to factors of interest may be removed. Nuanced future work will need to address this as past work has noted relations between MR image quality and general cognition, body mass index, and clinical group status [6, 8, 38].

Of note, there are many important limitations of our data and our results that must be highlighted. First, the public access dataset we used here, the Healthy Brain Network, is not a truly random sample. The dataset has a limited age range (5-21 years of age) and also employed a community-referred recruitment model. Study advertisements are specifically targeting families who have concerns about one or more psychiatric symptoms in their child. Given these factors, it is perhaps not surprising that our human raters excluded a large number of MRI scans. The Healthy Brain Network scanned many individuals who would not typically be involved with MRI research (e.g., youth with high levels of psychopathology and other development challenges), and therefore perhaps less likely to produce high quality data. However, the data loss rate seen in our project is actually in keeping with reports from past groups [39, 40]. Research teams interested in neurodevelopment and working in pediatric samples may think about use of prospective motion correction tools that localize the head position throughout the scan [41–43]. Second, Freesurfer is only one approach to deriving measures from structural MRI scans. Other metrics, such as voxel-based morphometry or region of interest drawing, may be similarly impacted by image quality. These approaches, however, often depend on tissue segmentation and would likely also be influenced by image quality. This should be investigated in the future by research teams employing such methods. Finally, we used a composite measure of image quality, constructed in the CAT12 toolbox. This may be influencing some of the results reported. There are many metrics of image quality, each potentially capturing unique aspects of noise relevant for MRI morphometry. We relied on this aggregated metric that combined noise to contrast ratio, coefficient of joint variation, inhomogeneity to contrast ratio, and root mean squared voxel resolution.

Expanding on this last issue, how to measure image quality is an area of much needed research. Here, being able to have a single “*grade*” (output by CAT12) motivated our decision to use this toolbox. We believe that researchers working in applied disciplines could use this single metric in their work to do quality control assessments, as well as a potential control variable in statistical models. Studies in the future could take an integrated approach to different measures connecting automated metrics of image quality (i.e., CAT12, Freesurfer’s Euler Number, MRIQC, Qoala-T) to trained human ratings and “crowd sourced” judgements of MR images [44, 45]. Such future work will need to balance how to reduce down these multiple metrics to fewer variables (to aid applied research teams) while isolating unique sources of noise. We feel that CAT12 is a reasonable starting point, as it is quick to run (∼18 minutes/subject), has a relatively easy to use interface, and does not require intense computational resources.

## 6. CONCLUSIONS

Limitations, notwithstanding, we demonstrate that direct measures of structural imaging quality are strongly linked to commonly-used structural MRI measures, as well as participant age. Importantly, we show that variations in image quality are strongly related to derivation of brain anatomy. Accounting for variations in image quality could impact results from applied studies (focused on age, clinical status, etc.). Unique to the work, we used more direct measures of structural MRI quality rather than proxies of motion and noise. In the future, research groups may consider accounting for such measures in analyses focused on individual differences in age, cognitive functioning, psychopathology, and other factors. This may lead to greater reproducibility in reported effects, as well as a way to minimize any potential spurious associations.

## Supporting information

Supplemental Analyses Examining MRI Quality and Different Subsamples of Project

## Declarations

## Availability of data and material

Relevant code and data is available at: https://github.com/jlhanson5/Gilmore_Buser_Hanson_CAT12_Freesurfer

## Authors’ contributions

Ms. Gilmore complete initial analyses, while Mr. Buser completed data processing. Ms. Gilmore wrote an initial draft of the manuscript. Dr. Hanson completed additional analyses and revised drafts of the manuscript.

