## Supplemental Analyses Examining MRI Quality and Different Subsamples of Project for "Variations in Structural MRI Quality Significantly Impact Commonly-Used Measures of Brain Anatomy"

**S1***.* **Additional Information about Assessment of General Cognitive Ability**

The Full-Scale IQ is derived from 7 of the 10 primary subtests: two Verbal Comprehension subtests, one Visual Spatial subtest, two Fluid Reasoning subtests, one Working Memory subtest, and one Processing Speed subtest. Verbal Comprehension and Fluid Reasoning are weighted more heavily in the Full-Scale IQ to reflect the importance of crystallized and fluid abilities in modern intelligence models (Wechsler, 2014).

**S2A. Quality Control Training Protocol**

Visual inspection of T1w Structural MRI scans was performed using the fslview program from the FSL neuroimaging package. Each subject was sequentially opened in the program and visualized across multiple slices in the axial, coronal, and sagittal view before an assessment score was decided on.

Research assistants were trained with textual descriptions and example screenshots to recognize a set of common artefacts: ghosting, blur, susceptibility, truncation, zipper, and spikes. Ghosting was described as “a vague impression or copy resembling the primary image but shifted.” Blur was described with analogy to blurred photographs: as an artefact relating to less clarity in anatomical features or divisions. Susceptibility was described as non-localized distortions in signal intensity” caused by the presence of some matter possessing metallic properties and often manifesting as a dark spot around the metallic matter (signal dropout) and an excessively bright area just beyond it (signal pileup). Truncation, as mild amounts of ring-like bands of brighter signal which occur close to tissue boundaries. Zipper artefacts were described as one or more lines of static on the frequency axis, as caused by electromagnetic interference from external sources. The final sort, spiking, was described as a wave-like pattern of signal loss across the brain, which affects the image globally and at a constant frequency.

After a period of training and familiarization with fslview and the common types of artefacts, research assistants were each given a wave of subjects to visually assess and score from 1-10, with brief descriptions of the perceived issue. The reviewers were informed that the scoring scale passed subjects with scores at or above a 6 and an automated column in excel reflected the pass/fail outcome for each subject based on the inputted score. Exemplar MR images of a score of 1, a 5, and 9.5 are shown in Figure S1. Figure S2 shows different example participant MRI scans with average scores, percentage of our original sample (N=388) that was included with ratings at (or above) these scores, and the percentage of Freesurfer cortical outputs related to image quality (as assessed by CAT12).

**S2B. Quality Control Scoring and Processing**

For each wave, scores were collected from six raters. Across these undergraduate research assistants, ratings were reasonably distributed and reasonably correlated. In terms of distributions, density plots for all of raters are shown in Figure S3A. The mean of all ratings of all raters was 6.145 (+/- 2.298) and the median was 6. Of note, one rater [“*Rater 5*”] did not rate 21 (out 388; ~5%) scans due to human error (i.e., omission from rating folder).

Ratings were well-correlated (bivariate correlations between raters, ranging from 0.794-0.921, as shown also in Figure S3B. Analysis of intraclass correlations (ICCs) indicated a strong level of reliability for these ratings. ICCs for absolute agreement were ICC(A,1) = 0.819 (95%-Confidence Interval: 0.779 <ICC<0.851); for consistency [average], reliability statistics were ICC(C,6) = 0.969 (95%-Confidence Interval: 0.964<ICC<0.974). A histogram of the average ratings is shown in Figure S4. Each score was z-scored by subtracting that rater’s average score and dividing it by the standard deviation for that rater. After z-scoring the scores were averaged across raters and those that were greater than the cutoff point was kept. The cutoff was also calculated through the same z-scoring mechanism of taking a score of 6.0, z-scoring it with respect to each rater, and taking the average z-score across the raters.

**S3. Analyses Examining Passing Visual Inspect, with CAT12 and Euler Number**

Motivated by past work using Freesurfer’s Euler Number in relation to cortical thickness, we also probed connections between visual quality control checks (scan inclusion / exclusion) and this MRI scan quality metric. Similar to CAT12, Euler number was a significant predictor in a logistic regression model (p<.005, with the dependent binary variable of include/exclude and Euler number as the independent variable). Twenty-one participants were unable to be processed in Freesurfer due to errors in that software package (i.e., poor quality scans causing issues in different processing steps). Of note, CAT12 and Euler Number were strongly correlated (r=-0.904, p<.0005; as shown in Figure S5).

In a logistic regression model with both CAT12 and Euler number, both are significant predictors of scan inclusion and exclusion (both p<.008, DV: Scan Inclusions, IVs: Euler Number and CAT12). Comparing the estimates output for this model, CAT12 appears to be a more robust predictor (z= 4.977, p<.005) compared to Euler Number (z=-2.614, p=.008). The difference between these z-scores (z_difference_=2.363) would suggest this is a statistically significant difference (p=.018 for a two-tailed test). Receiver operating characteristic curves showed similar performance between the three methods (1: Euler Number only; 2: CAT12 Ratings only; 3. Both Quality Control Variables in the logistic models; these are shown in Figure S6). All of these models had similar Areas under the curve (*Model 1*: 0.984; *Model 2*: 0.989; *Model 3*: 0.990).

**S4. Association Between CAT12 Scan Rating and Freesurfer Outputs, Controlling for Participant Age**

Given that participant age was significantly related to CAT12 quality scores, these associations may be influencing connections between CAT12 scan rating and Freesurfer outputs. To these ends, and similar to analyses reported in the main manuscript, we, therefore, examined correlations between structural MRI quality and 158 morphometric outputs from Freesurfer (68 mean cortical surface area estimates from the DK atlas; 68 mean cortical thickness estimates also from the DK atlas; 22 subcortical regions); however, in these analyses, we constructed linear regression models examining Freesurfer outputs (area, thickness, or subcortical volume) as the dependent variable, and age and CAT12 score as the independent variables.

For these models, and related to cortical surface area, there was variability in how image quality related to mean surface area from differ brain parcels (T-statistic range=-1.004-4.734). In aggregate, this association was modest (Mean t-statistic=1.424+/- 1.255); however, in ten areas, the association between image quality and mean surface area was significant (p _fdr-corrected_<.05), even after correcting for multiple comparisons (as shown in Table S1 and Figure S7). All relations were positive in nature, with higher quality relating to higher surface area in different brain parcels. For cortical thickness, there was again variability in relation between mean thickness for parcels and image quality (T-statistic range=-1.535-6.671), with modest associations in the aggregate (Mean t-statistic=2.474 +/- 2.038). However, relations between image quality and cortical thickness for thirty-seven regions was significant (p _fdr-corrected_<.05), even after correcting for multiple comparisons (as shown in Table S2 and Figure S8). Again, all relations were positive in nature, with higher quality relating to higher thickness in different brain parcels. Finally, for subcortical volume, similar patterns were seen (t-statistic range=-0.974-2.807; mean t-statistic=0.897 +/- 0.964, as shown in Table S3 and Figure S9). Of note, no regions were related to image quality (p _fdr-corrected_<.05) after correcting for multiple comparisons.

**S5. Association Between CAT12 Scan Rating and Freesurfer Outputs, Controlling for Freesurfer’s Euler Number**

Given correlations between CAT12 quality scores and Freesurfer’s Euler number, we also completed exploratory analyses, similar to the main manuscript, examining correlations between structural MRI quality and 158 morphometric outputs from Freesurfer, but while controlling for Euler number. These analyses involved of linear regression models examining Freesurfer outputs (area, thickness, or subcortical volume) as the dependent variable, and Euler number and CAT12 score as the independent variables.

For these models, and related to cortical surface area, there was variability in how image quality related to mean surface area from differ brain parcels (t-statistic range=-1.082-3.262). In aggregate, this association was modest (Mean t-statistic=1.256+/- 1.060); however, in four areas, the association between image quality and mean surface area was significant (p _fdr-corrected_<.05), even after correcting for multiple comparisons (as shown in Table S4 and Figure S10). All relations were positive in nature, with higher quality relating to higher surface area in different brain parcels. For cortical thickness, there was again variability in relation between mean thickness for parcels and image quality (t-statistic range=-1.571 5.806), with modest associations in the aggregate (Mean t-statistic=2.689 +/- 1.667). However, relations between image quality and cortical thickness for forty-two regions was significant (p _fdr-corrected_<.05), even after correcting for multiple comparisons (as shown in Table S4 and Figure S11). Again, all relations were positive in nature, with higher quality relating to higher thickness in different brain parcels. Finally, for subcortical volume, similar patterns were seen (t-statistic range= -0.680-2.732; mean t-statistic=0.607 +/- 0.8328, as shown in Table S6 and Figure S12). Of note, no regions were related to image quality (p _fdr-corrected_<.05) after correcting for multiple comparisons.

**S6. Association Between CAT12 Scan Rating and Freesurfer Outputs, Only in Participants Excluded After Visual Quality Control by Human Raters**

To more deeply understand the impact of visual quality control by raters, we finally mirrored the analyses in our main manuscript, but this time only examining participants who failed visual quality control by raters; this was in contrast to the main manuscript where we only examined associations between MRI quality and Freesurfer outputs in those subjects that passed visual quality control by raters. This was again done in 158 morphometric outputs from Freesurfer. For these models, and related to cortical surface area, there was variability in how image quality related to mean surface area from differ brain parcels (t-statistic range= 3.97-13.06). In aggregate, this association was fairly large (Mean t-statistic= 8.535 +/- 2.094). The association between image quality and mean surface area was significant (p _fdr-corrected_<.05) for all 68 regions examined even after correcting for multiple comparisons (as shown in Table S7 and Figure S13). All relations were positive in nature, with higher quality relating to higher surface area in different brain parcels. For cortical thickness, there was again variability in relation between mean thickness for parcels and image quality (t-statistic range-2.034-12.72), with modest associations in the aggregate (Mean t-statistic=2.719+/- 2.129). However, relations between image quality and cortical thickness for forty-four regions was significant (p _fdr-corrected_<.05), even after correcting for multiple comparisons (as shown in Table S8 and Figure S14). Again, all relations were positive in nature, with higher quality relating to higher thickness in different brain parcels. Finally, for subcortical volume, similar patterns were seen (t-statistic range=-2.123- 7.491; mean t-statistic=2.708 +/- 2.891, as shown in Table S9 and Figure S15). Twelve regions were related to image quality (p _fdr-corrected_<.05) after correcting for multiple comparisons in these subjects that were failed visual quality checks.

**S7. Association Between Freesurfer Outputs and Freesurfer’s Euler Number, Controlling CAT12 Scan Rating**

Completing the analyses detailed in Section S5 above, we also completed exploratory analyses examining correlations between Freesurfer’s Euler Number and 158 morphometric outputs from Freesurfer, but when controlling for CAT12 quality score. These analyses involved of linear regression models examining Freesurfer outputs (area, thickness, or subcortical volume) as the dependent variable, and Euler number and CAT12 score as the independent variables. For these models, and related to cortical surface area, there was variability in how image quality related to mean surface area from differ brain parcels (t-statistic range=-2.496-2.691). In aggregate, this association was modest (Mean t-statistic=0.204 +/- 1.068). This association between Euler number and mean surface area was not significant for any brain areas when controlling for multiple comparisons (and accounting for CAT12 scores, shown in Table S10 and Figure S16). For cortical thickness, there was again variability in relation between mean thickness for parcels and image quality (t-statistic range=-1.422-5.846), with modest associations in the aggregate (Mean t-statistic=2.359 +/- 1.682). However, relations between image quality and cortical thickness for thirty-eight regions was significant (p _fdr-corrected_<.05), even after correcting for multiple comparisons (as shown in Table S11 and Figure S17). Again, all relations were positive in nature, with higher quality relating to higher thickness in different brain parcels. Finally, for subcortical volume, similar patterns were seen (t-statistic range= -2.159-0.837; mean t-statistic=-0.613 +/- 0.893, as shown in Table S12 and Figure S18). However, no regions were related to image quality (p _fdr-corrected_<.05) after correcting for multiple comparisons.

**S8. Open Science / Data Availability:**

Relevant analytic code is available at the following GitHub repository: <https://github.com/jlhanson5/Gilmore_Buser_Hanson_CAT12_Freesurfer>. This repo contains RMarkdown for reproducible statistical analyses, as well as quality control reports (as pdfs) output from the CAT12 toolbox. The structural MRI data used in this work is also public-access and available from: <http://fcon_1000.projects.nitrc.org/indi/cmi_healthy_brain_network/sharing_neuro.html>.

Table S1. Relations Between Cortical Surface Area and Structural MRI Quality (assessed by the CAT12 Toolbox), Controlling for Participant Age

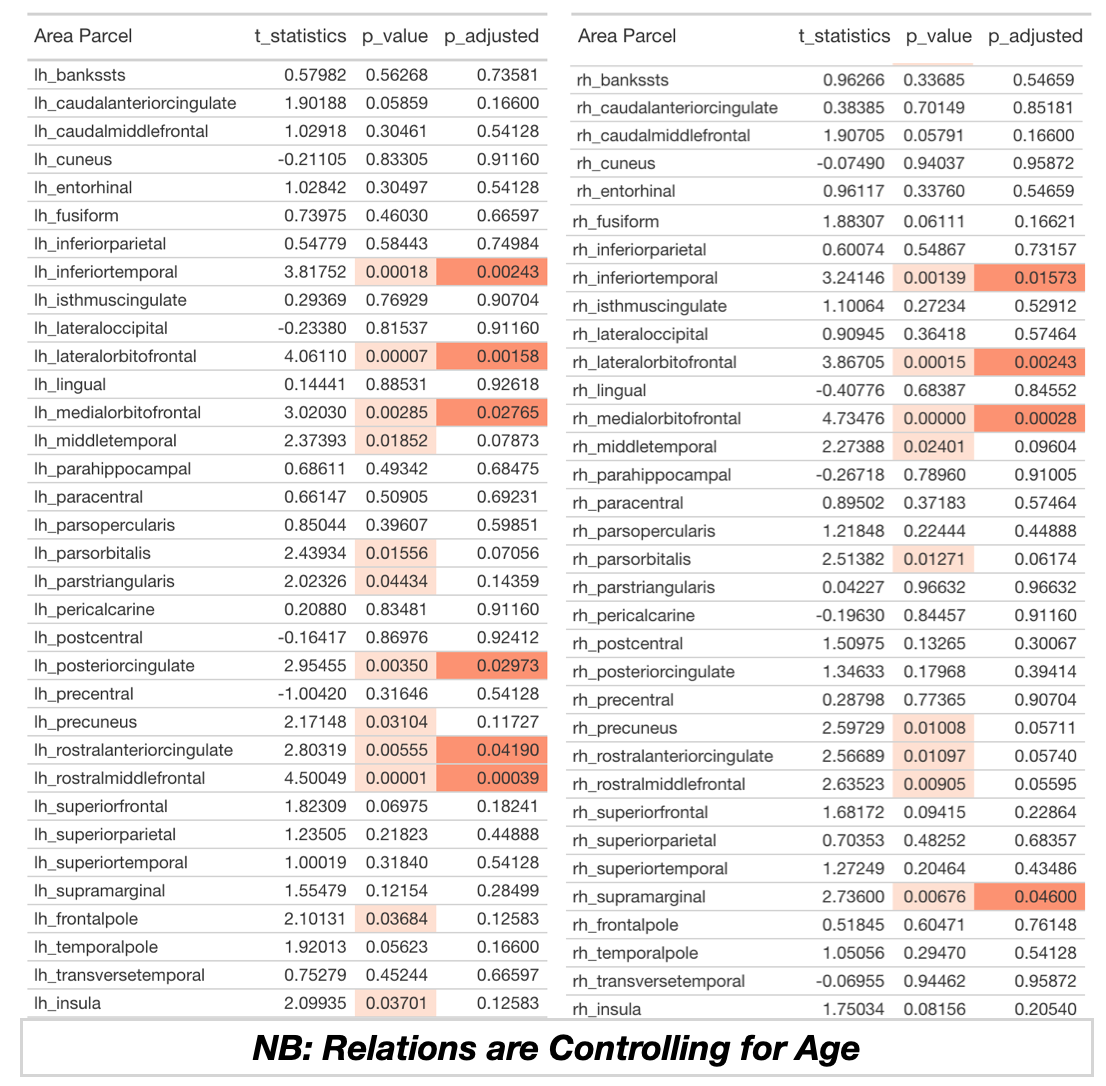

Caption: Table displays relations between MRI quality (CAT12 score) and cortical surface area for different brain parcels in Freesurfer’s DK atlas. In contrast to the results in the main manuscript, these analyses control for participant age. The left side of the table shows regions in the left hemisphere, while the right side shows the right hemisphere. On each side, region is in the first column, and t-statistic (of CAT12 and cortical thickness) is in the second column. The third column is the uncorrected p-value, while the fourth column is this test statistic corrected for multiple comparisons (for all 68 cortical parcels). Light orange highlighting indicates regions that were p<.05 (uncorrected), while darker orange highlighting indicates regions that were p<.05 (FDR corrected).

Table S2. Relations Between Cortical Thickness and Structural MRI Quality (assessed by the CAT12 Toolbox), Controlling for Participant Age

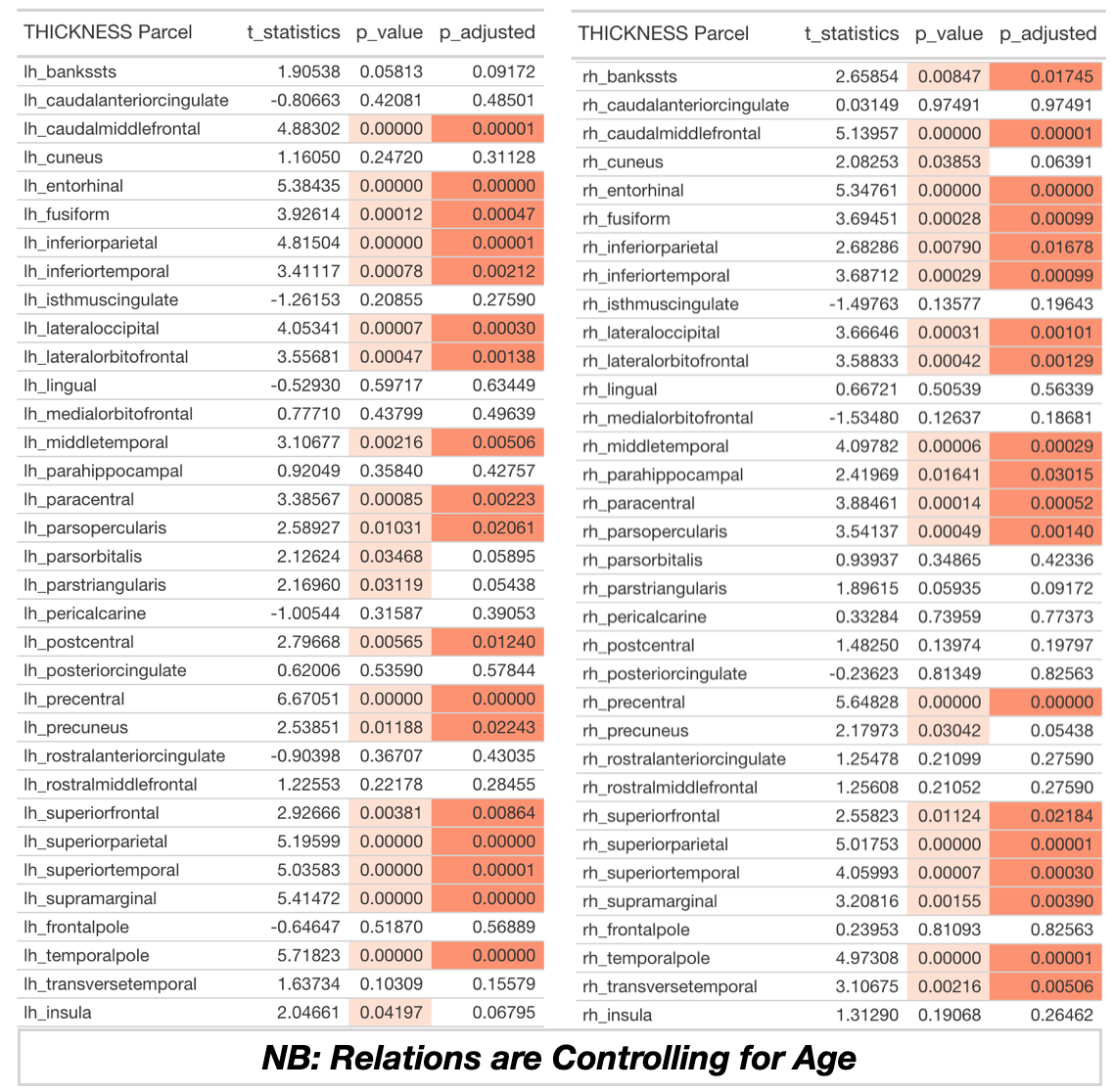

Caption: Table displays relations between MRI quality (CAT12 score) and cortical thickness for different brain parcels in Freesurfer’s DK atlas. In contrast to the results in the main manuscript, these analyses control for participant age. The left side of the table shows regions in the left hemisphere, while the right side shows the right hemisphere. On each side, region is in the first column, and t-statistic (of CAT12 and cortical thickness) is in the second column. The third column is the uncorrected p-value, while the fourth column is this test statistic corrected for multiple comparisons (for all 68 cortical parcels). Light orange highlighting indicates regions that were p<.05 (uncorrected), while darker orange highlighting indicates regions that were p<.05 (FDR corrected).

Table S3. Relations Between Subcortical Volumes and Structural MRI Quality (assessed by the CAT12 Toolbox), Controlling for Participant Age

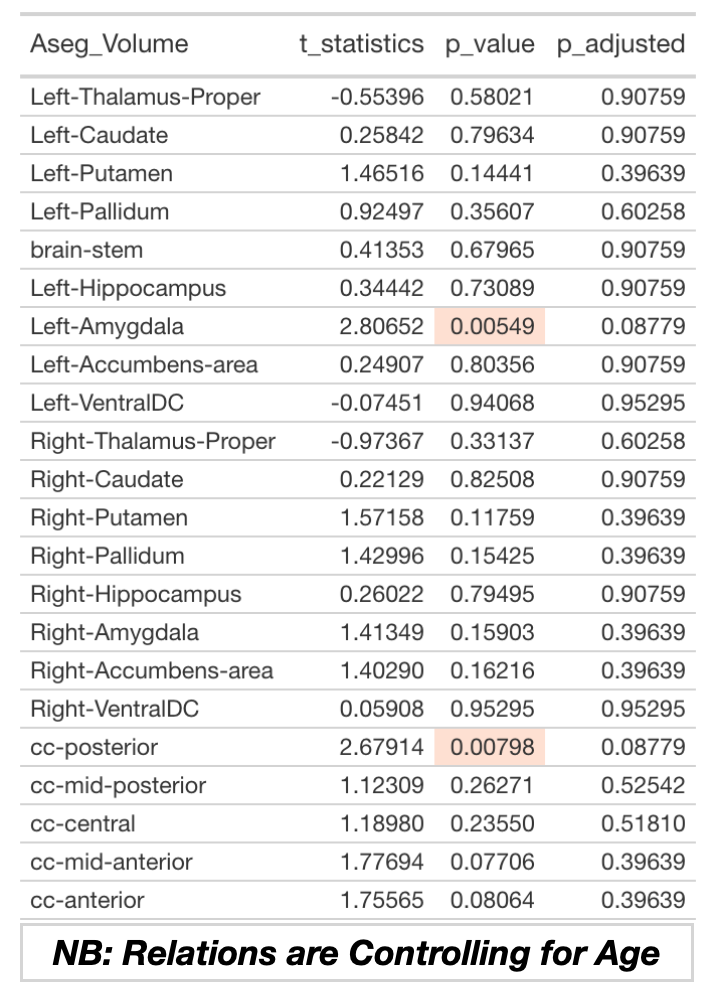

Caption: Table displays relations between MRI quality (CAT12 score) and subcortical volumes in Freesurfer’s ASEG atlas. In contrast to the results in the main manuscript, these analyses control for participant age. Region is in the first column, and t-statistic (of CAT12 and subcortical volume) is in the second column. The third column is the uncorrected p-value, while the fourth column is this test statistic corrected for multiple comparisons (for all 68 cortical surface comparisons). Light orange highlighting indicates regions that were p<.05 (uncorrected), while darker orange highlighting indicates regions that were p<.05 (FDR corrected).

Table S4. Relations Between Cortical Surface Area and Structural MRI Quality (assessed by the CAT12 Toolbox), Controlling for Freesurfer’s Euler Number

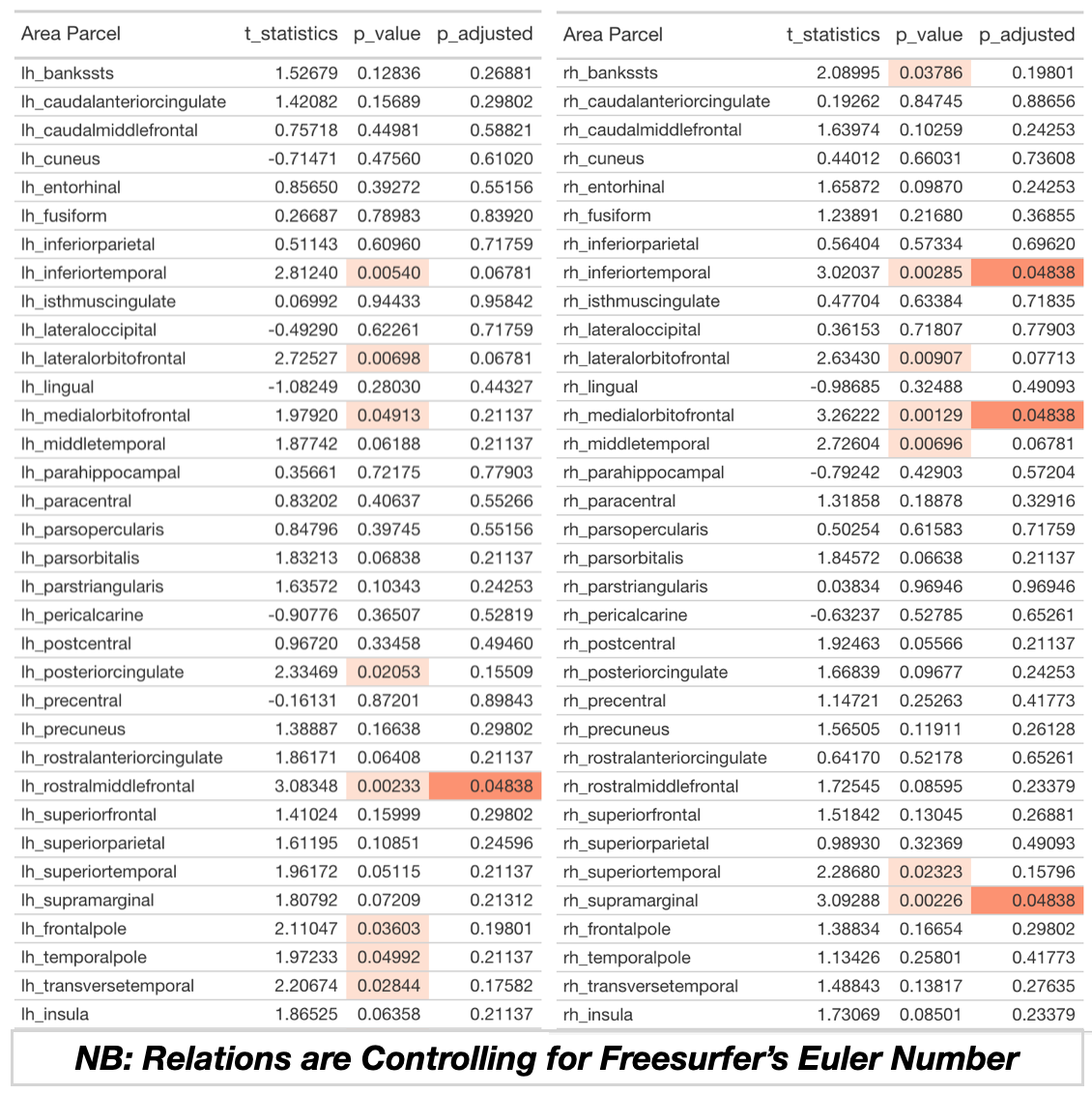

Caption: Caption: Table displays relations between MRI quality (CAT12 score) and cortical surface area for different brain parcels in Freesurfer’s DK atlas. In contrast to the results in the main manuscript, these analyses control for Freesurfer’s Euler Number. The left side of the table shows regions in the left hemisphere, while the right side shows the right hemisphere. On each side, region is in the first column, and t-statistic (of CAT12 and cortical thickness) is in the second column. The third column is the uncorrected p-value, while the fourth column is this test statistic corrected for multiple comparisons (for all 68 cortical parcels). Light orange highlighting indicates regions that were p<.05 (uncorrected), while darker orange highlighting indicates regions that were p<.05 (FDR corrected).

Table S5. Relations Between Cortical Thickness and Structural MRI Quality (assessed by the CAT12 Toolbox), Controlling for Freesurfer’s Euler Number

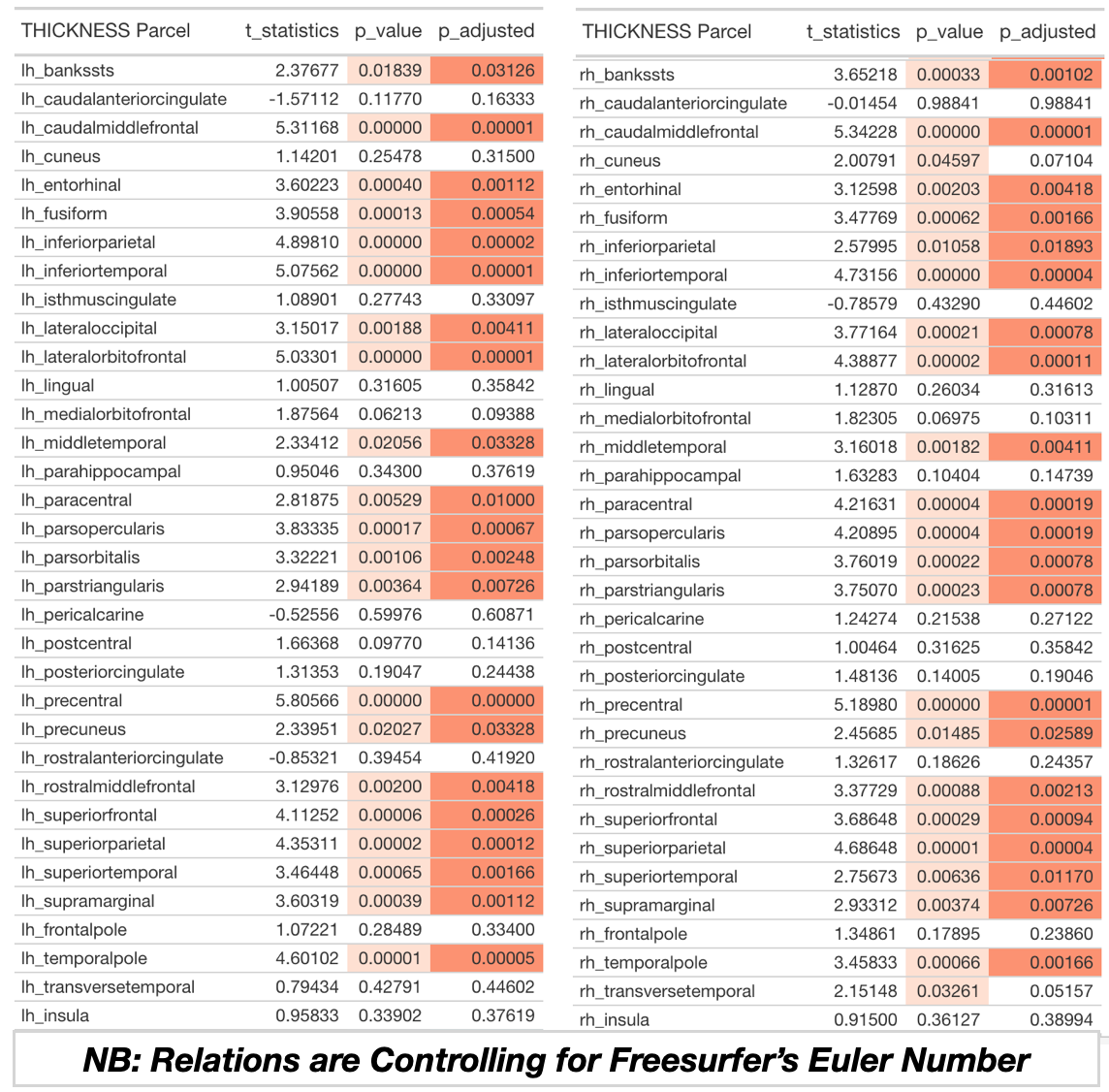

Caption: Caption: Table displays relations between MRI quality (CAT12 score) and cortical thickness for different brain parcels in Freesurfer’s DK atlas. In contrast to the results in the main manuscript, these analyses control for Freesurfer’s Euler Number. The left side of the table shows regions in the left hemisphere, while the right side shows the right hemisphere. On each side, region is in the first column, and t-statistic (of CAT12 and cortical thickness) is in the second column. The third column is the uncorrected p-value, while the fourth column is this test statistic corrected for multiple comparisons (for all 68 cortical parcels). Light orange highlighting indicates regions that were p<.05 (uncorrected), while darker orange highlighting indicates regions that were p<.05 (FDR corrected).

Table S6. Relations Between Subcortical Volumes and Structural MRI Quality (assessed by the CAT12 Toolbox), Controlling for Freesurfer’s Euler Number

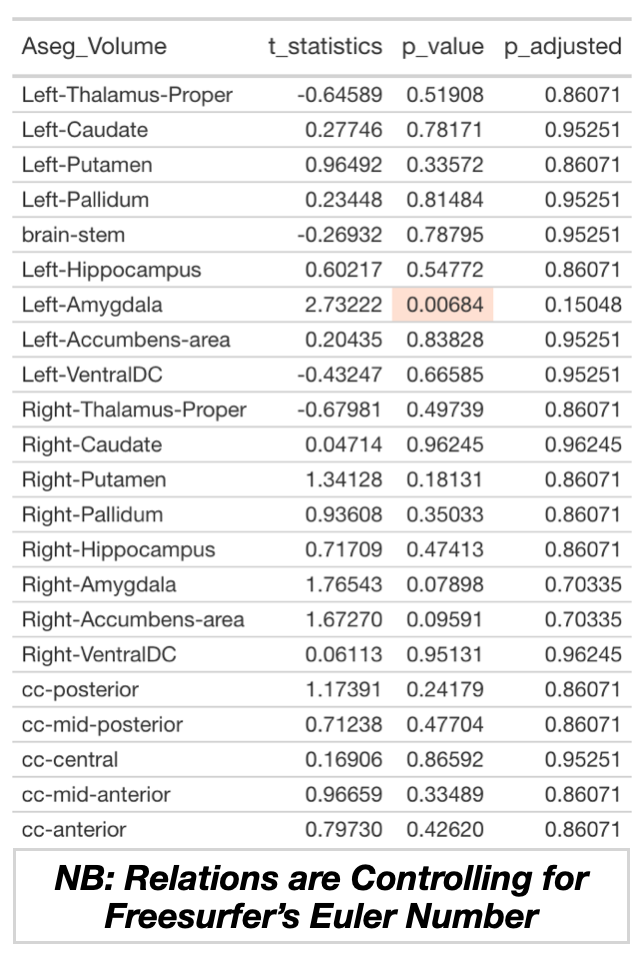

Caption: Table displays relations between MRI quality (CAT12 score) and subcortical volumes in Freesurfer’s ASEG atlas. In contrast to the results in the main manuscript, these analyses control for Freesurfer’s Euler Number. Region is in the first column, and t-statistic (of CAT12 and subcortical volume) is in the second column. The third column is the uncorrected p-value, while the fourth column is this test statistic corrected for multiple comparisons (for all 68 cortical surface comparisons). Light orange highlighting indicates regions that were p<.05 (uncorrected), while darker orange highlighting indicates regions that were p<.05 (FDR corrected).

Table S7. Relations Between Cortical Surface Area and Structural MRI Quality (assessed by the CAT12 Toolbox) in Participants that Failed Visual Quality Checks

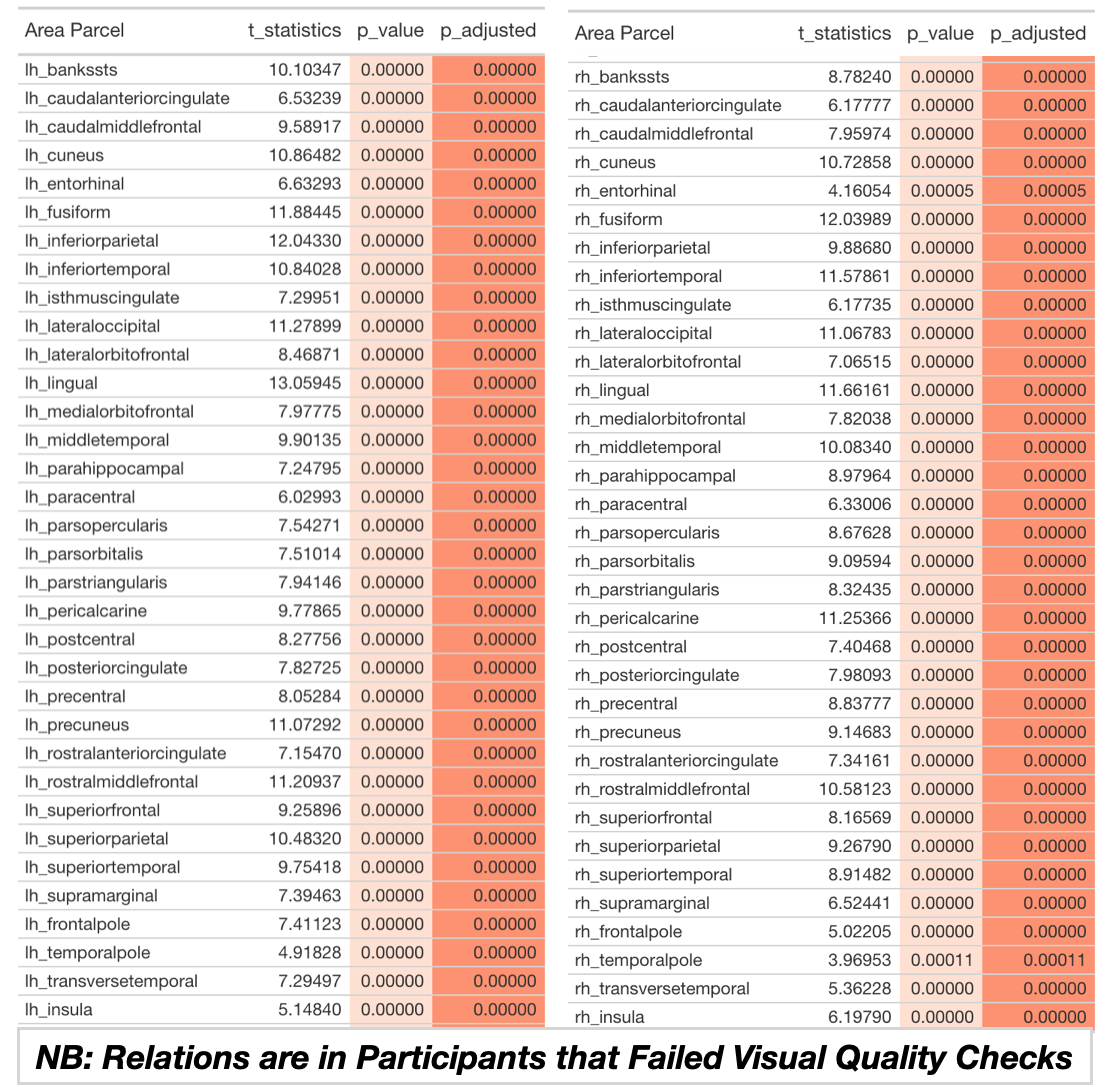

Caption: Caption: Table displays relations between MRI quality (CAT12 score) and cortical surface area for different brain parcels in Freesurfer’s DK atlas. In contrast to the results in the main manuscript, these analyses are for participants who failed visual quality checks (by human raters). The left side of the table shows regions in the left hemisphere, while the right side shows the right hemisphere. On each side, region is in the first column, and t-statistic (of CAT12 and cortical thickness) is in the second column. The third column is the uncorrected p-value, while the fourth column is this test statistic corrected for multiple comparisons (for all 68 cortical parcels). Light orange highlighting indicates regions that were p<.05 (uncorrected), while darker orange highlighting indicates regions that were p<.05 (FDR corrected).

Table S8. Relations Between Cortical Thickness and Structural MRI Quality (assessed by the CAT12 Toolbox) in Participants that Failed Visual Quality Checks

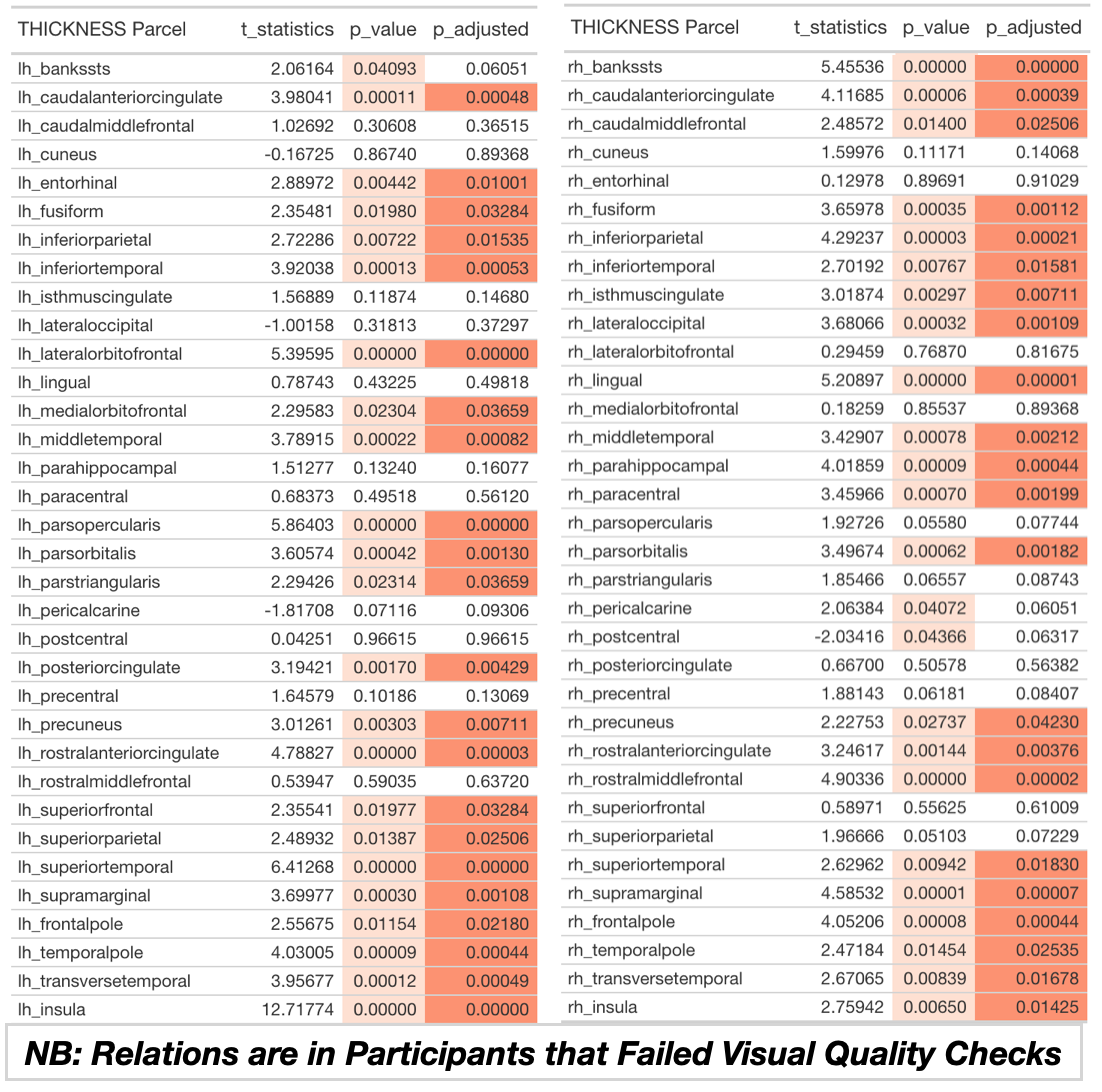

Caption: Caption: Table displays relations between MRI quality (CAT12 score) and cortical thickness for different brain parcels in Freesurfer’s DK atlas. In contrast to the results in the main manuscript, these analyses are for participants who failed visual quality checks (by human raters). The left side of the table shows regions in the left hemisphere, while the right side shows the right hemisphere. On each side, region is in the first column, and t-statistic (of CAT12 and cortical thickness) is in the second column. The third column is the uncorrected p-value, while the fourth column is this test statistic corrected for multiple comparisons (for all 68 cortical parcels). Light orange highlighting indicates regions that were p<.05 (uncorrected), while darker orange highlighting indicates regions that were p<.05 (FDR corrected).

Table S9. Relations Between Subcortical Volumes and Structural MRI Quality in Participants that Failed (Human) Visual Quality Control Checks

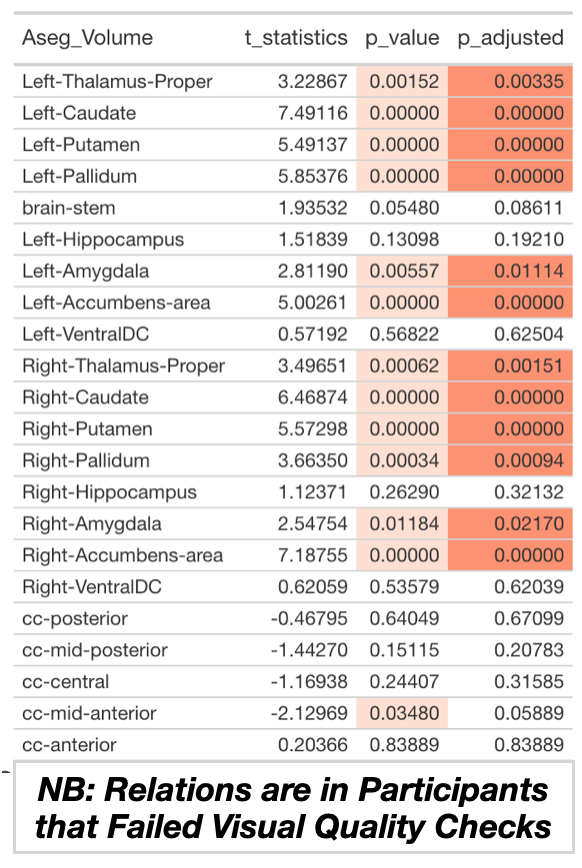

Caption: Table displays relations between MRI quality (CAT12 score) and subcortical volumes in Freesurfer’s ASEG atlas. In contrast to the results in the main manuscript, these analyses are for participants who failed visual quality checks (by human raters). Region is in the first column, and t-statistic (of CAT12 and subcortical volume) is in the second column. The third column is the uncorrected p-value, while the fourth column is this test statistic corrected for multiple comparisons (for all 68 cortical surface comparisons). Light orange highlighting indicates regions that were p<.05 (uncorrected), while darker orange highlighting indicates regions that were p<.05 (FDR corrected).

Table S10. Relations Between Cortical Surface Area and Freesurfer’s Euler Number, Controlling for CAT12 Toolbox Quality Score

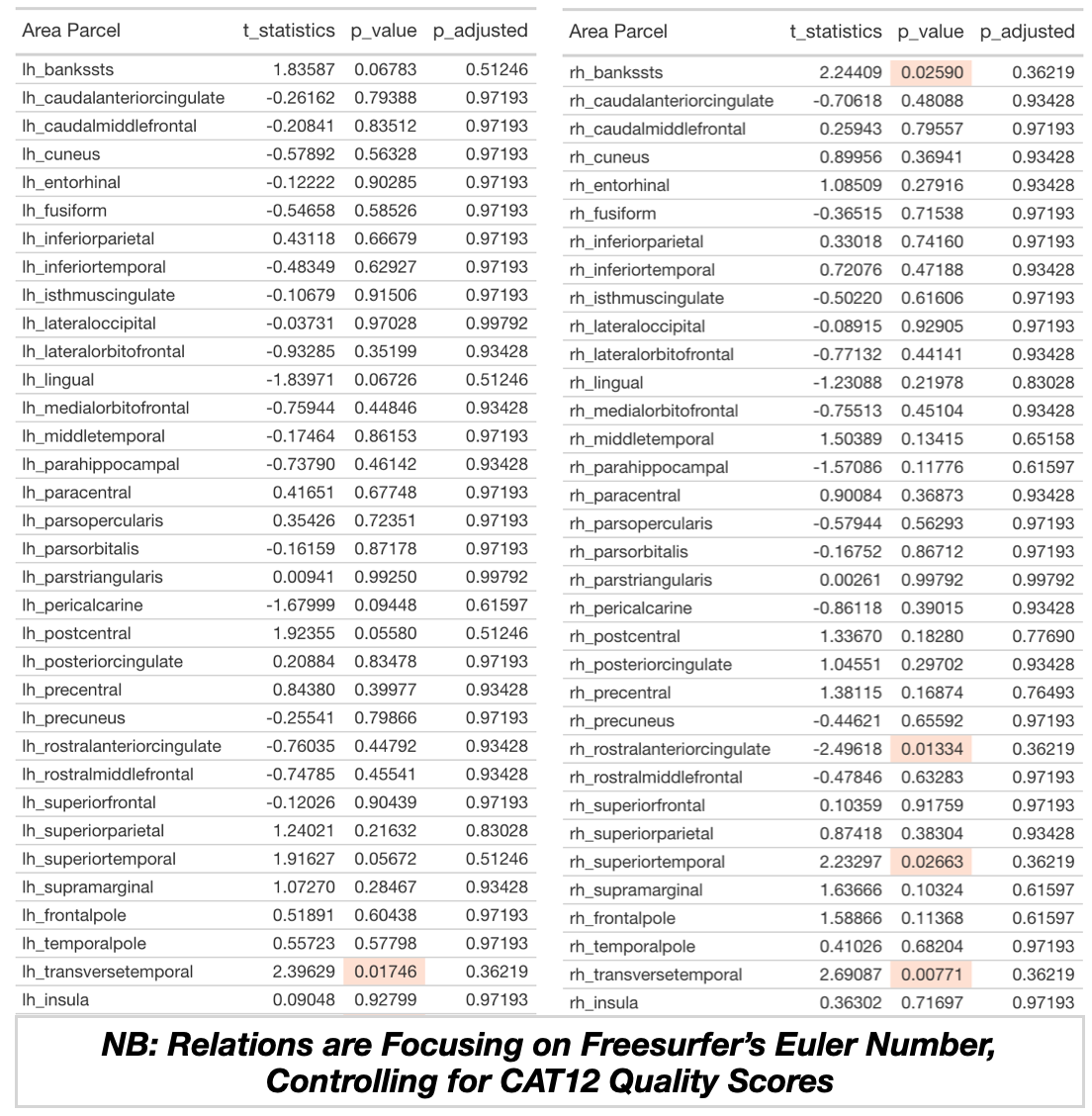

Caption: Caption: Table displays relations between Euler Number and cortical surface area for different brain parcels in Freesurfer’s DK atlas. In contrast to the results noted above, these analyses focus on Freesurfer’s Euler Number and control for CAT12 Quality Scores. The left side of the table shows regions in the left hemisphere, while the right side shows the right hemisphere. On each side, region is in the first column, and t-statistic (of CAT12 and cortical thickness) is in the second column. The third column is the uncorrected p-value, while the fourth column is this test statistic corrected for multiple comparisons (for all 68 cortical parcels). Light orange highlighting indicates regions that were p<.05 (uncorrected), while darker orange highlighting indicates regions that were p<.05 (FDR corrected).

Table S11. Relations Between Cortical Thickness and Freesurfer’s Euler Number, Controlling for CAT12 Toolbox Quality Score

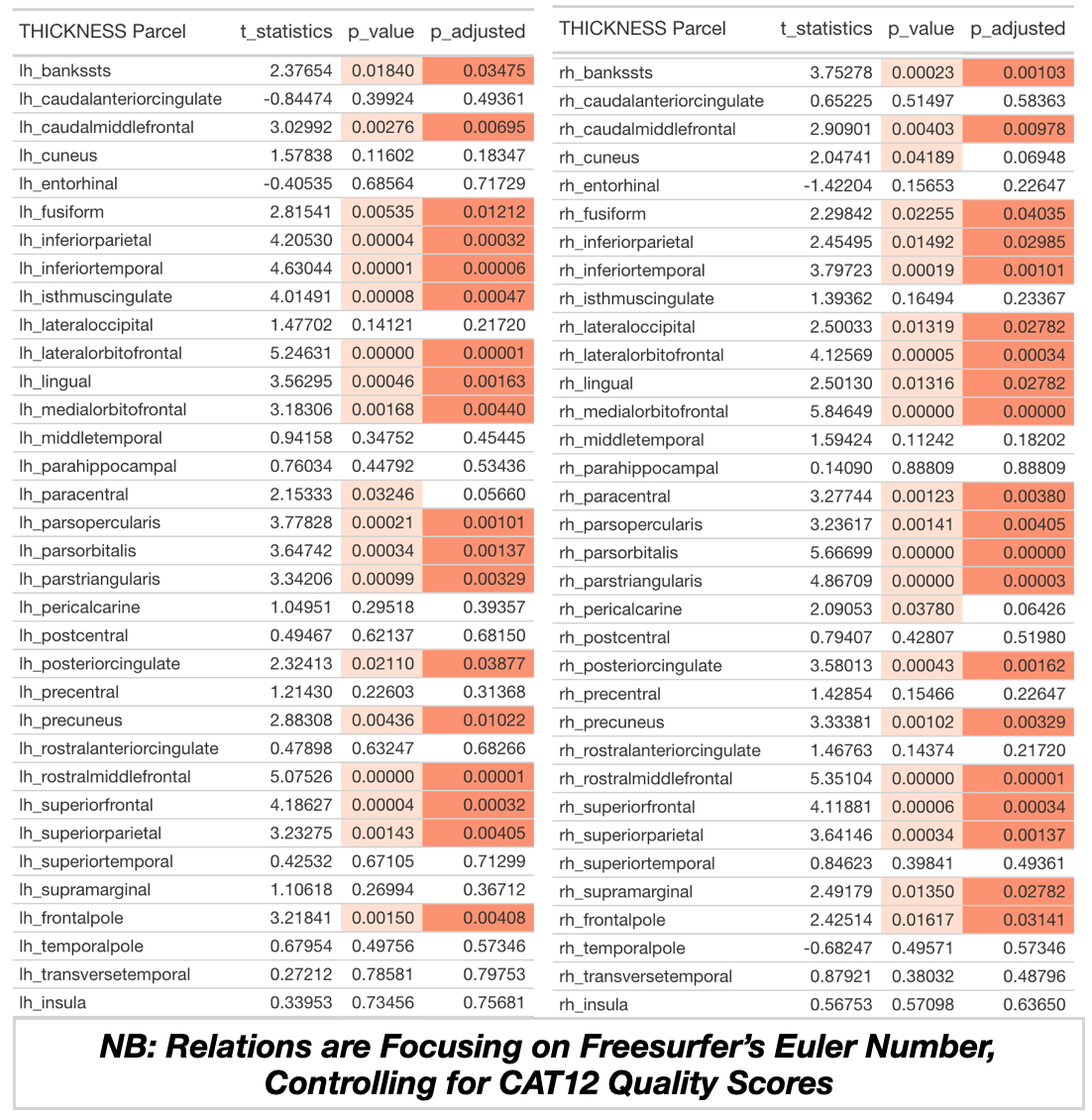

Caption: Table displays relations between Euler Number and cortical thickness for different brain parcels in Freesurfer’s DK atlas. In contrast to the results noted above, these analyses focus on Freesurfer’s Euler Number and control for CAT12 Quality Scores. The left side of the table shows regions in the left hemisphere, while the right side shows the right hemisphere. On each side, region is in the first column, and t-statistic (of CAT12 and cortical thickness) is in the second column. The third column is the uncorrected p-value, while the fourth column is this test statistic corrected for multiple comparisons (for all 68 cortical parcels). Light orange highlighting indicates regions that were p<.05 (uncorrected), while darker orange highlighting indicates regions that were p<.05 (FDR corrected).

Table S12. Relations Between Subcortical Volumes and Euler Number, Controlling for CAT12 Scores

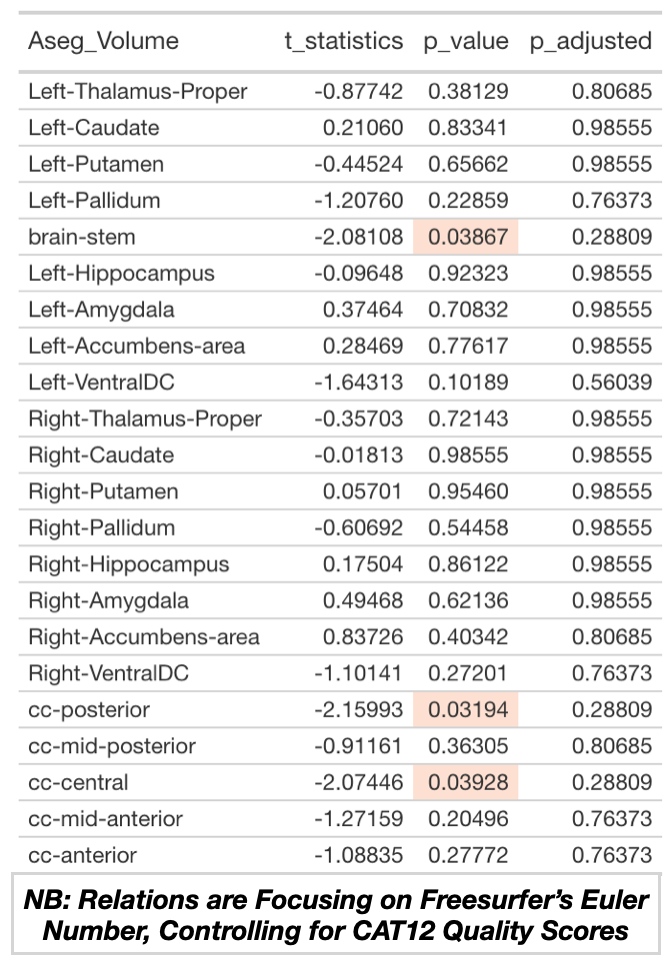

Caption: Table displays relations between Euler Number and subcortical volumes in Freesurfer’s ASEG atlas. In contrast to the results noted above, these analyses focus on Freesurfer’s Euler Number and control for CAT12 Quality Scores.Region is in the first column, and t-statistic (of CAT12 and subcortical volume) is in the second column. The third column is the uncorrected p-value, while the fourth column is this test statistic corrected for multiple comparisons (for all 68 cortical surface comparisons). Light orange highlighting indicates regions that were p<.05 (uncorrected), while darker orange highlighting indicates regions that were p<.05 (FDR corrected).

Figure S1.

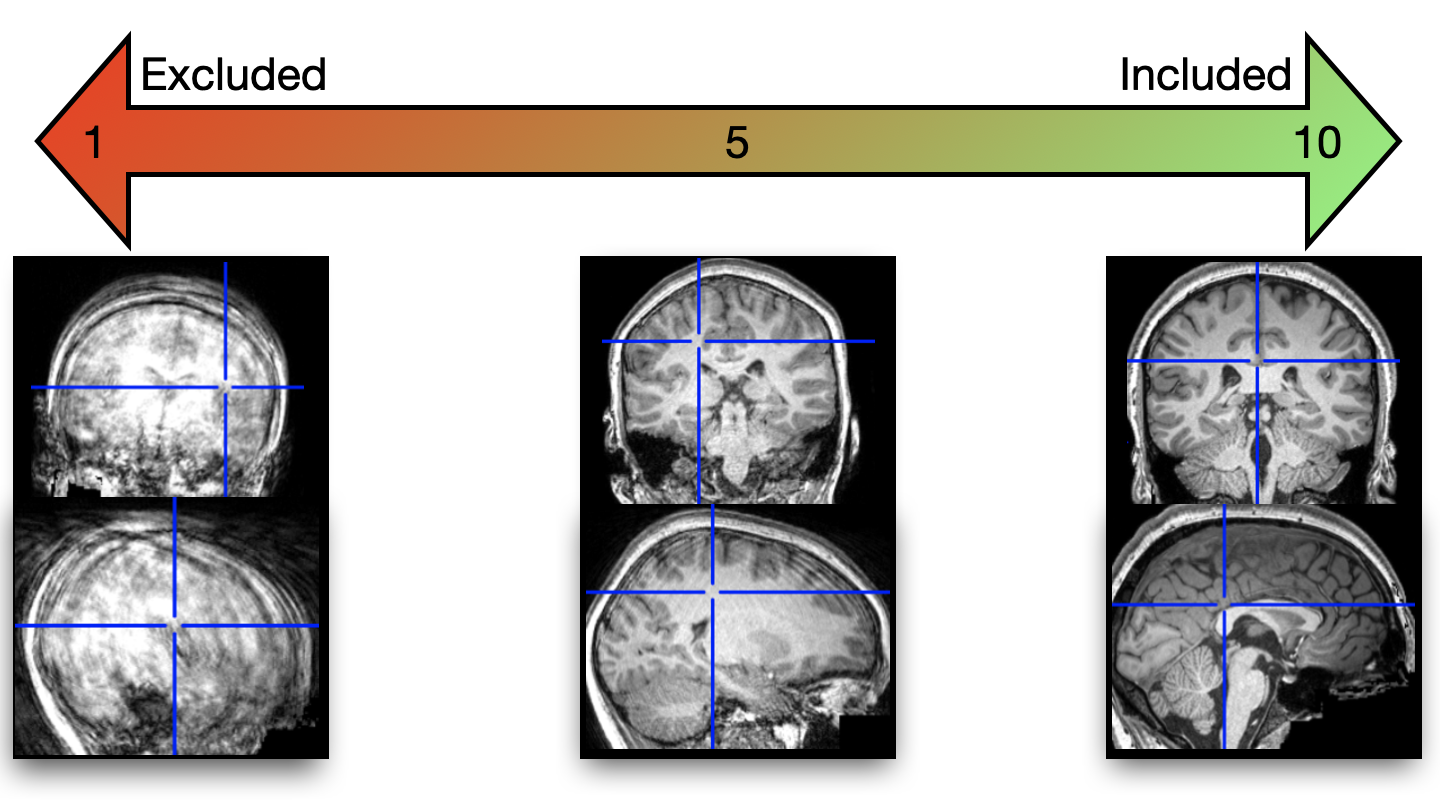

Caption: Graphic depiction of MRI and typical human visual rater scores. A scan with high-levels of motion is shown on the left side and received a score of 1. In the middle, a scan of reasonable quality received a score of 5. The sagittal image displays a great deal of ringing artifacts related to participant movement, while the coronal image’s artifacts are more salient in more superior portions of the cortex. On the right side, a higher quality scan that received a score of 9.5 is shown. This scan was one that would be retained in relevant analyses.

Table S2.

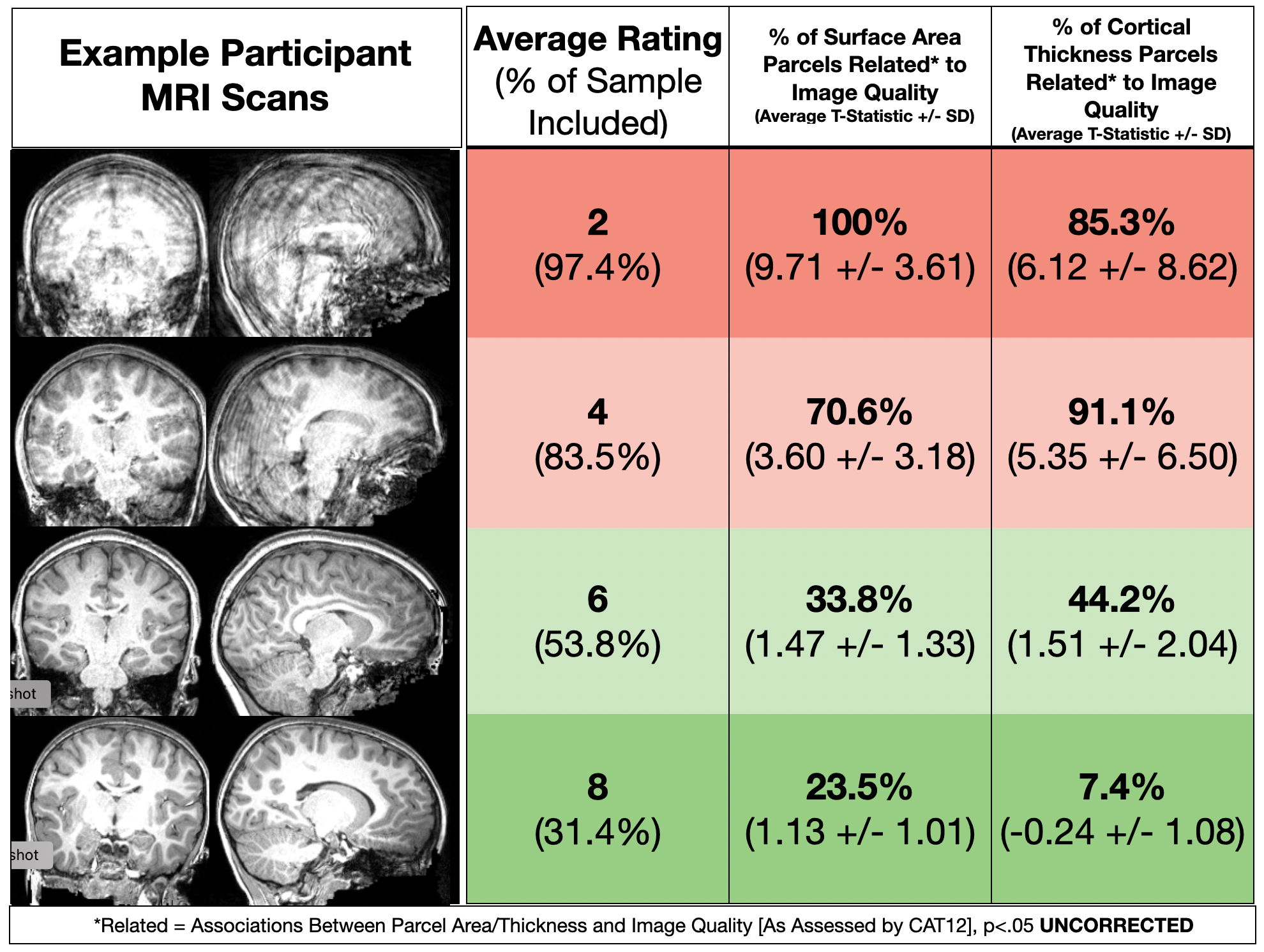

Caption: This graphic shows example participant MRI scans with average scores (left and middle of the figure) of 2, 4, 6, and 8 (going top to bottom). Percentage of the original sample that would be included at each score cutoff is also noted. On the right side of the image, the percentage of Freesurfer cortical outputs related to image quality (as assessed by CAT12) is shown. Of note, this is for p<0.05 **uncorrected** (and this is notable given that there are 68 comparisons for both surface area and cortical thickness). The average t-score and standard-deviation of relations between Freesurfer cortical outputs related to image quality is also noted).

Figure S3.

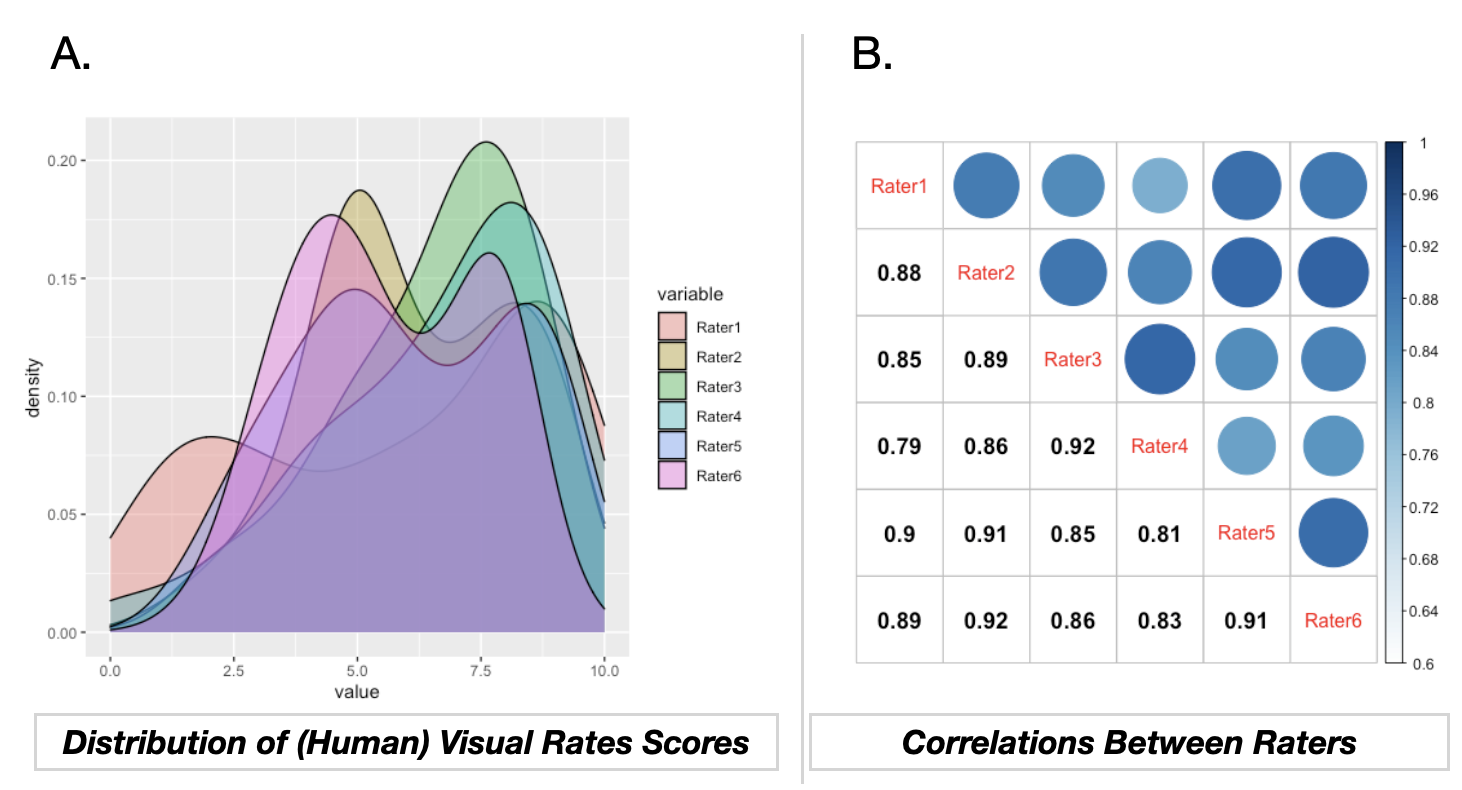

Caption: Panel A shows a density plot of our six research assistant ratings of MRI scans. Each rater is shown in a different color. Panel B shows the correlations between each rater scores (the bottom of that panel is the numeric correlation, while the top displays the same information as dots that vary in color and size based on the strength of the correlation).

Figure S4.

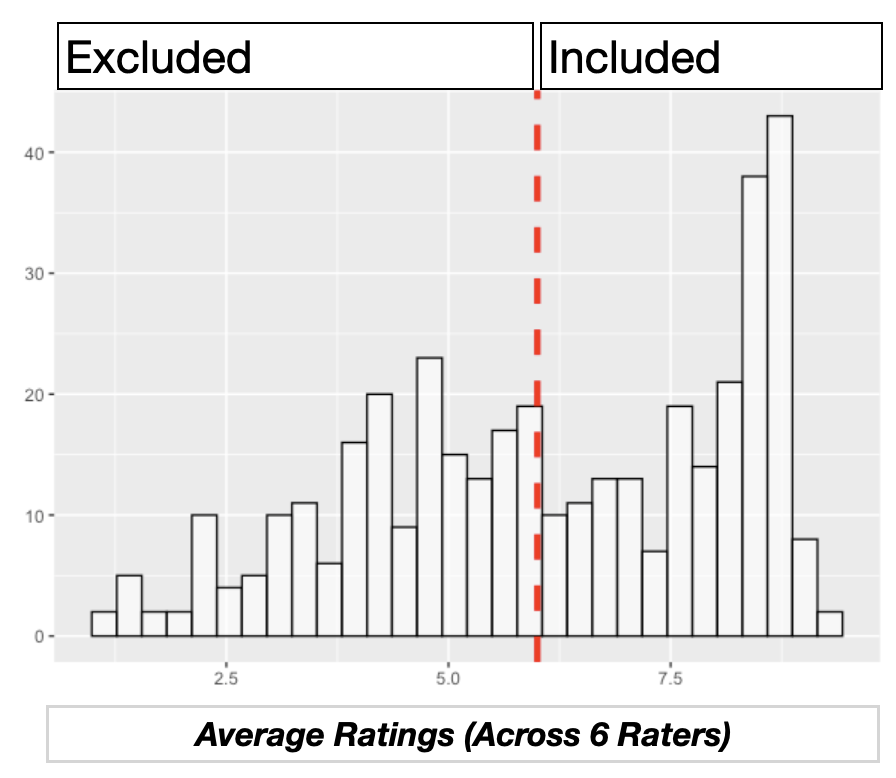

Caption: A histogram showing the distribution of average ratings (across our human raters). A red dotted line depicts our cutoff point of 6 (for scan inclusion).

Figure S5.

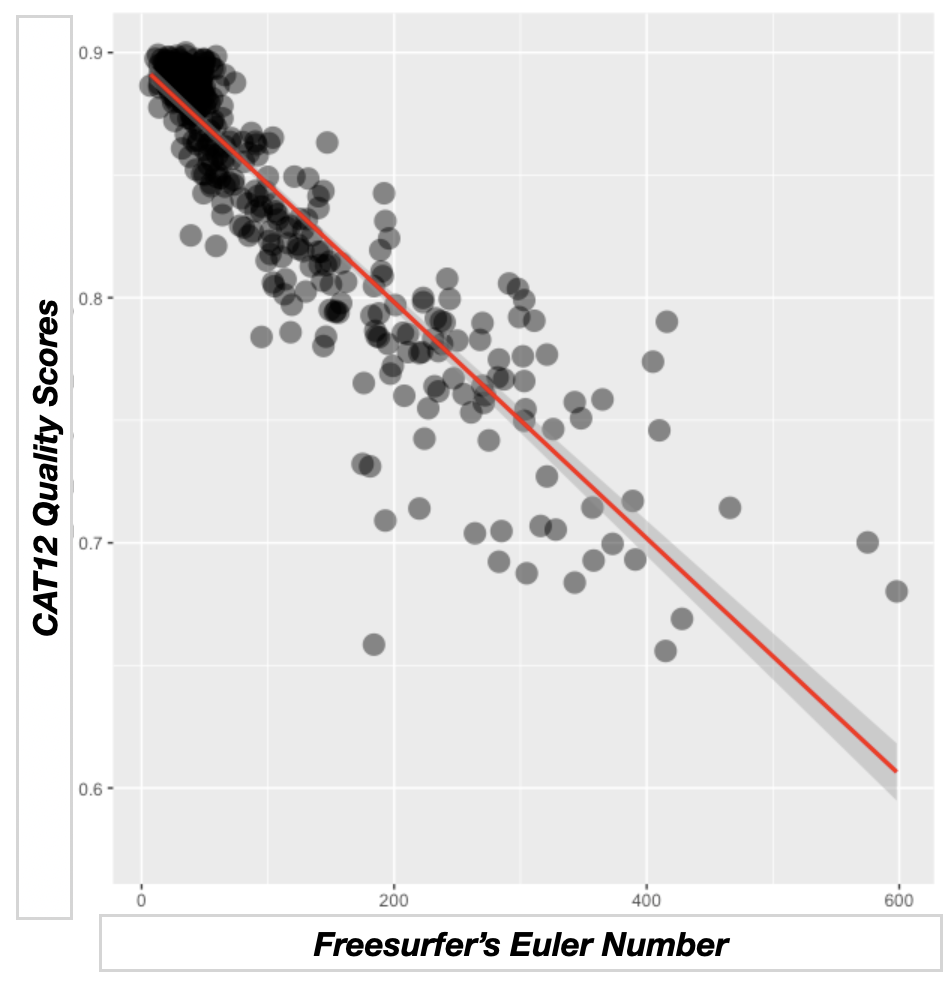

Caption: A scatterplot showing the relation between Freesurfer’s Euler number (on the horizontal axis) and CAT12 Quality Score (on the vertical axis). These variables were strongly correlated (r=-0.904, p<.0005).

Figure S6.

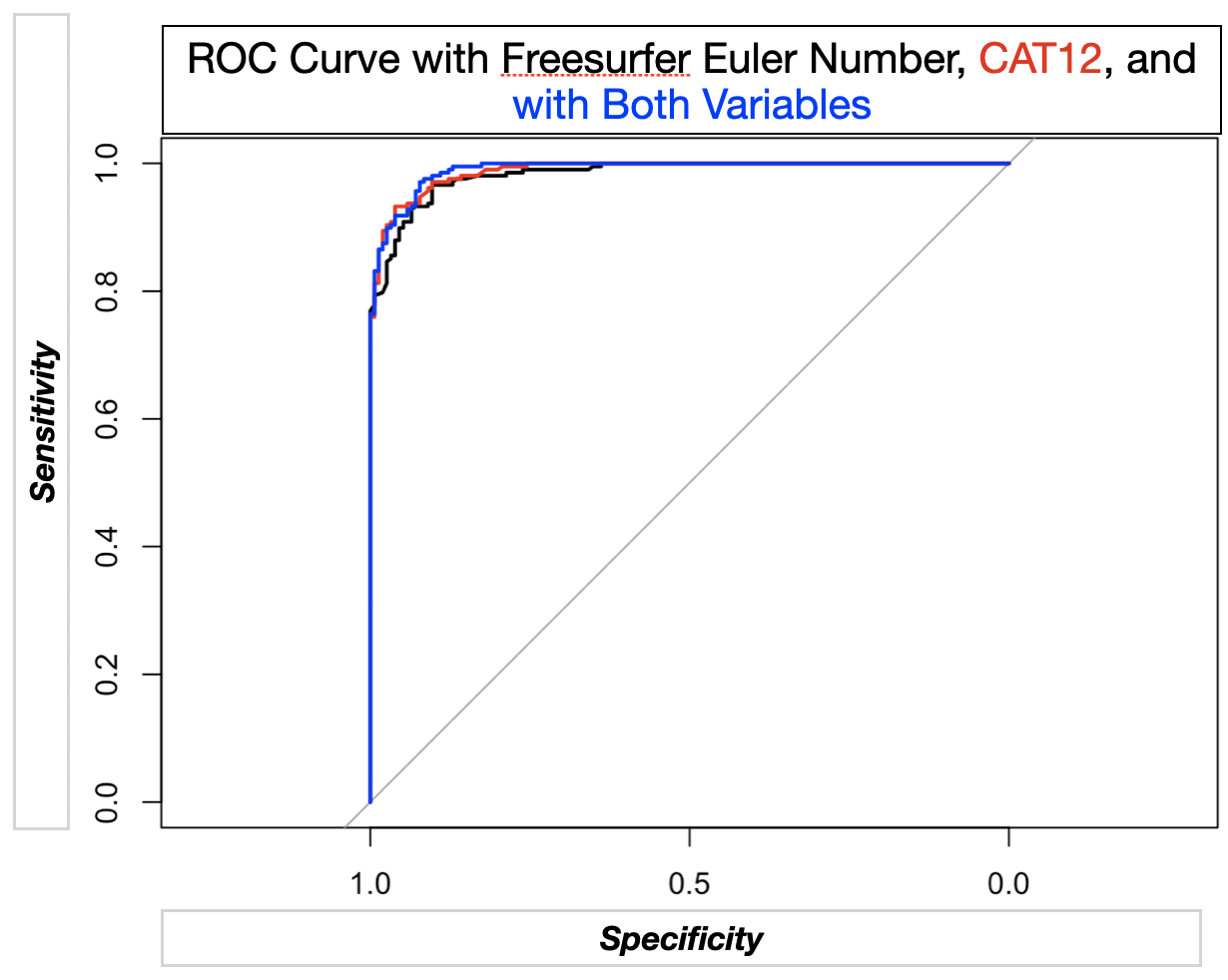

Caption: ROC curves showing sensitivity and specificity for Freesurfer’s Euler number (shown in black) and CAT12 (shown in red) in separate logistic regression models where the image quality metric (Euler number or CAT12 was the independent variable) and passing human rater visual checks of quality (as a binary, was the dependent variable). ROC curves for a logistic model containing both Euler number and CAT12 are shown in blue.

Figure S7.

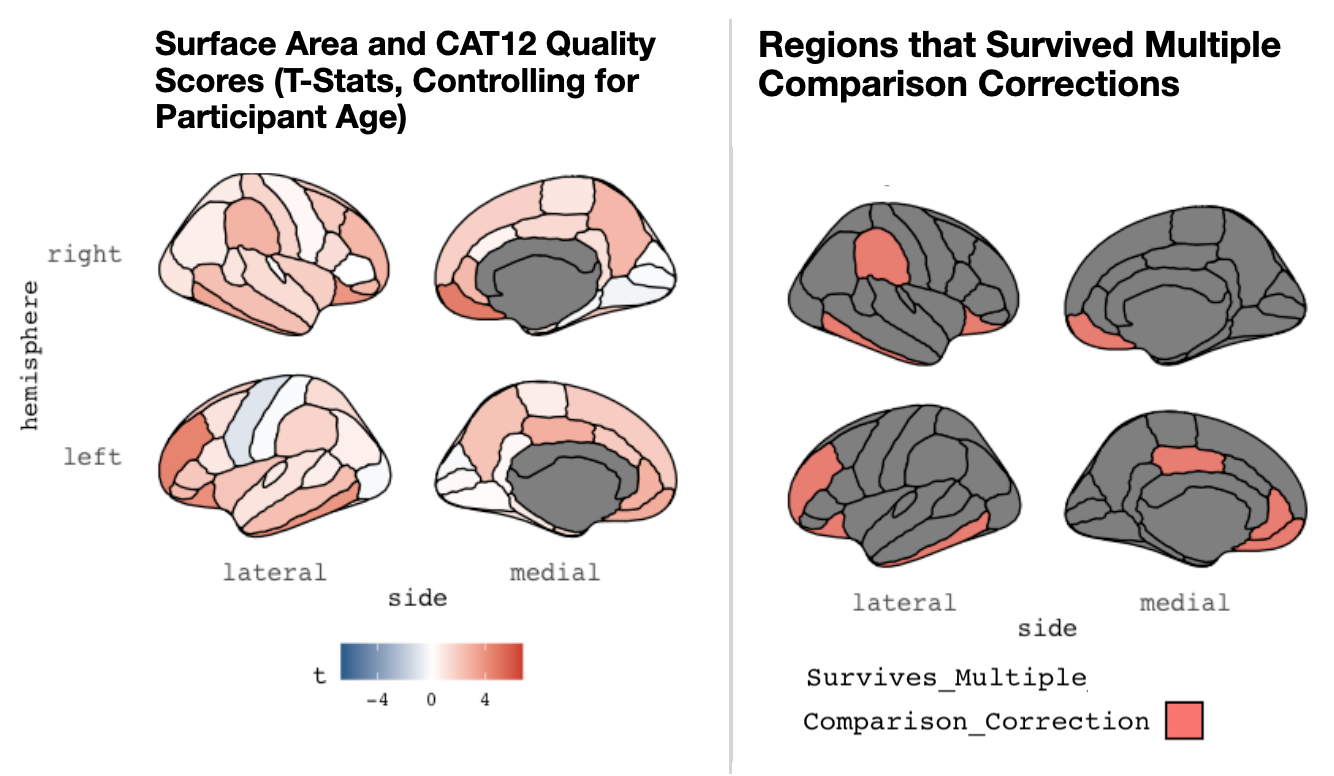

Caption: A graphic depiction (from the R library *ggseg*) showing associations between image quality (as assessed by the CAT12 toolbox) and derived (mean) cortical surface area. This is shown for the Desikan atlas commonly used in Freesurfer. These analyses controlled for participant age. Lateral and medial views are shown for the right (top) and left (bottom) hemispheres. The left panel shows the overall t-statistics for the relation in each parcel, while the right panel shows parcels where the relation between surface area and image quality survives multiple comparisons.

Figure S8.

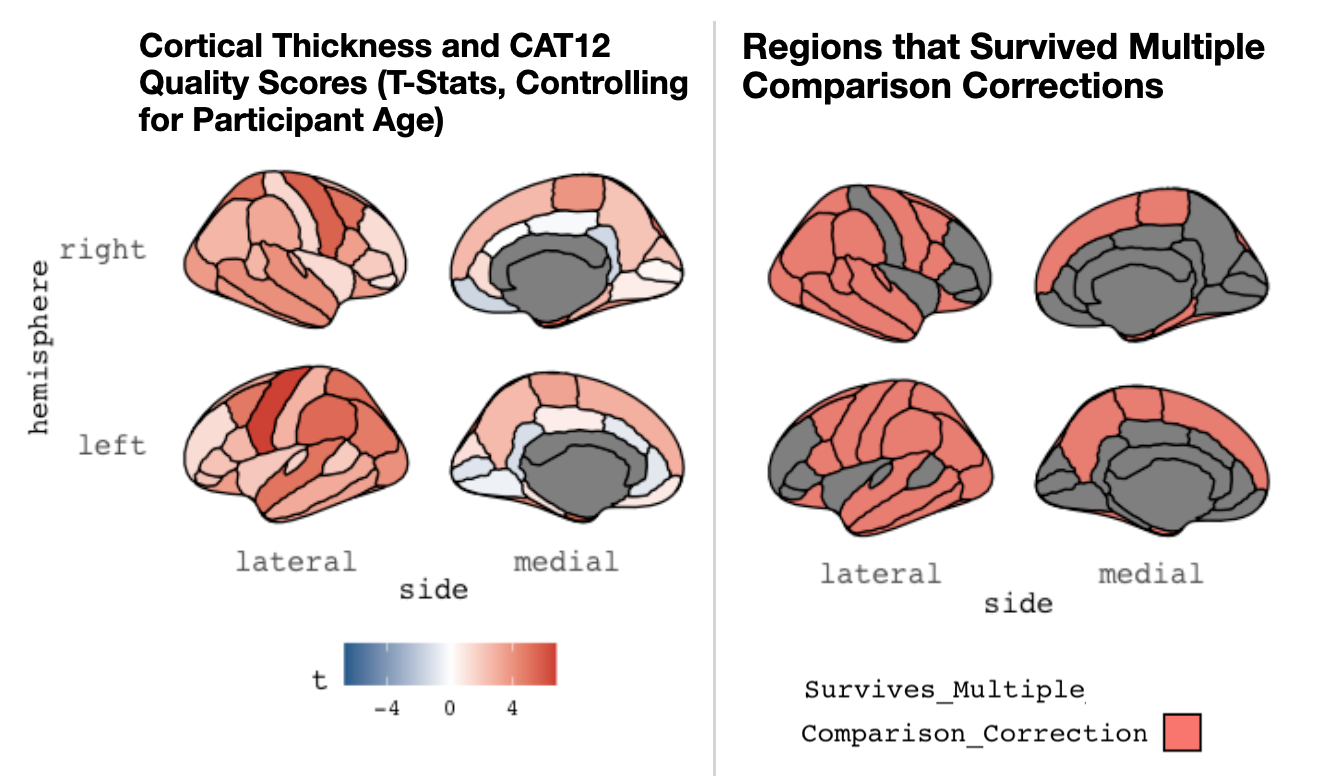

Caption: A graphic depiction (from the R library *ggseg*) showing associations between image quality (as assessed by the CAT12 toolbox) and derived (mean) cortical thickness. This is shown for the Desikan atlas commonly used in Freesurfer. These analyses controlled for participant age. Lateral and medial views are shown for the right (top) and left (bottom) hemispheres. The left panel shows the overall t-statistics for the relation in each parcel, while the right panel shows parcels where the relation between surface area and image quality survives multiple comparisons.

Figure S9.

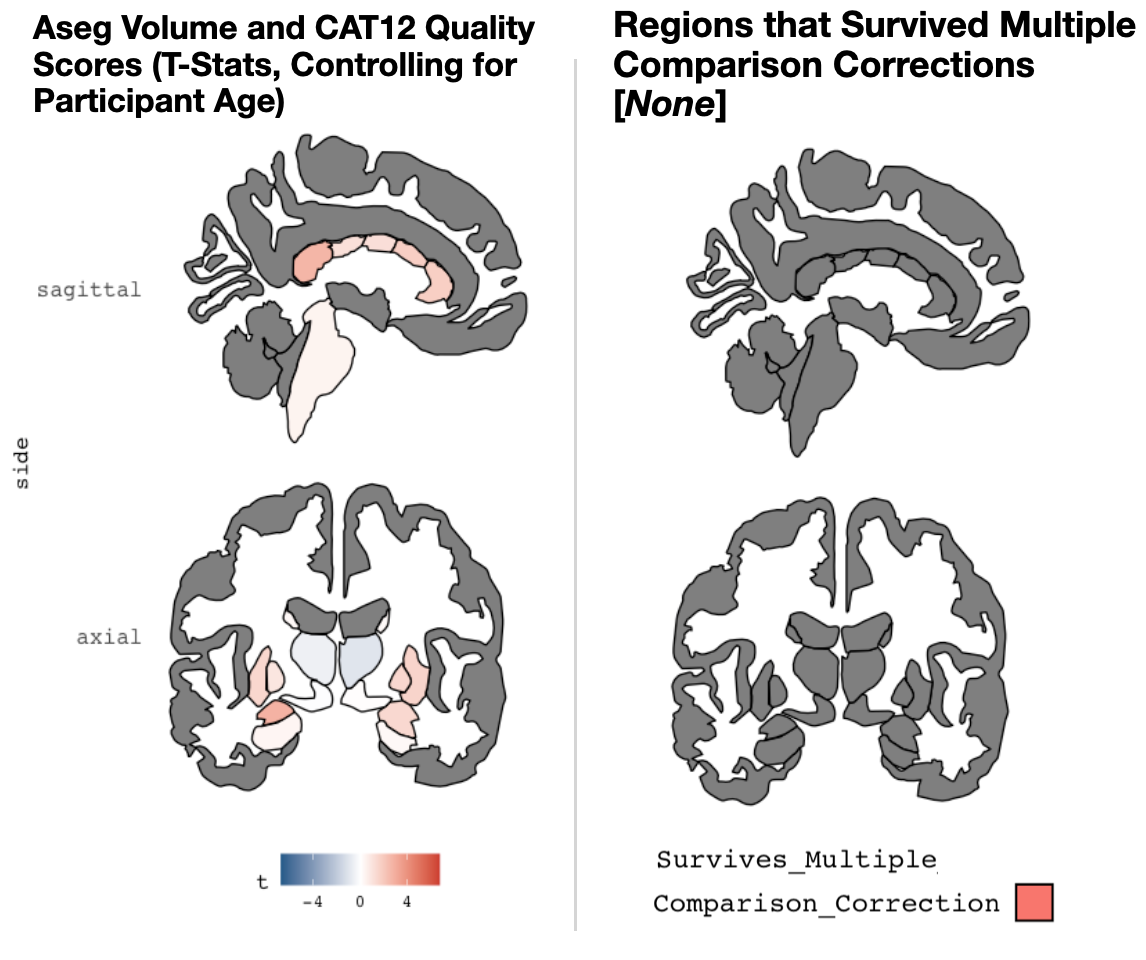

Caption: A graphic depiction (from the R library ggseg) showing associations between image quality (assessed by the CAT12 Toolbox) and subcortical volumes. This is shown for the Freesurfer ASEG atlas. Coronal (left) and sagittal (right) views are shown. These analyses controlled for participant age. The left panel shows the t-statistic for the relation in each subcortical volume, while the right panel shows parcels where the relation between volume and image quality survives multiple comparisons. Of note, no regions survived multiple comparisons correction.

Figure S10.

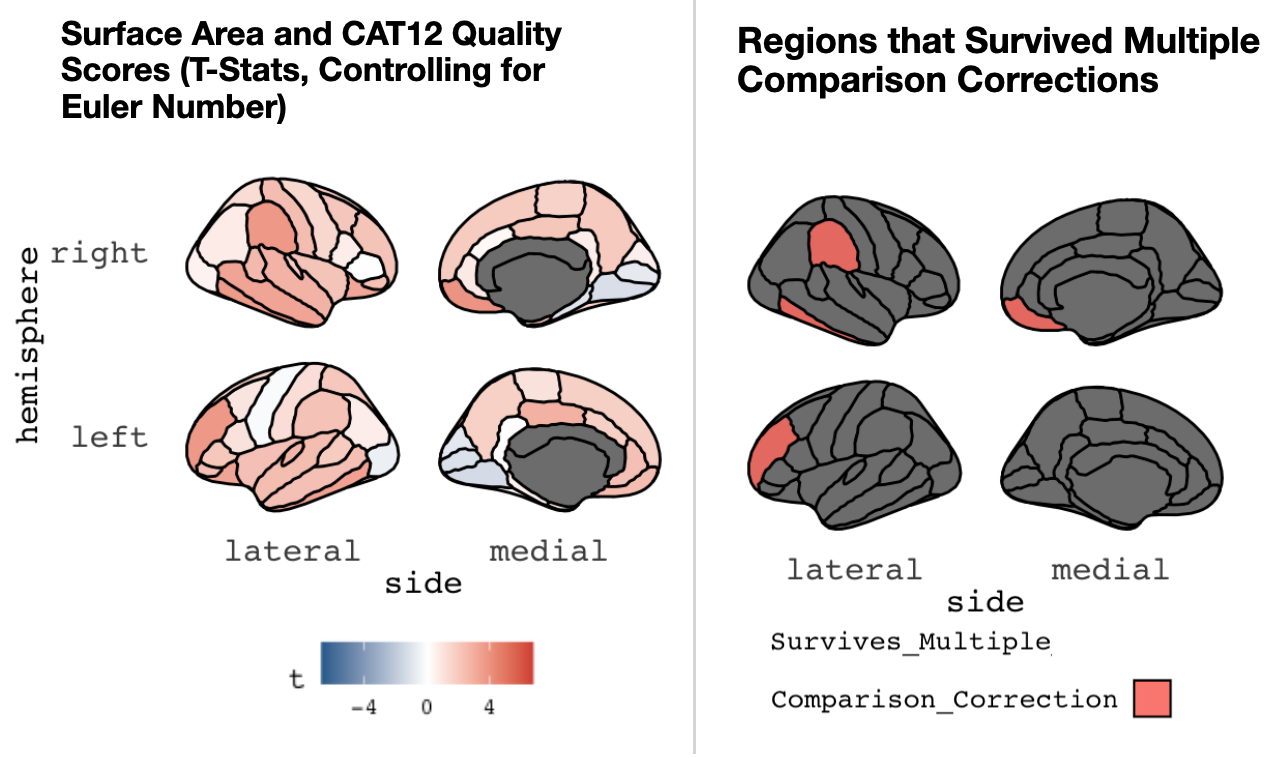

Caption: A graphic depiction (from the R library *ggseg*) showing associations between image quality (as assessed by the CAT12 toolbox) and derived (mean) cortical surface area. This is shown for the Desikan atlas commonly used in Freesurfer. These analyses controlled for Freesurfer Euler Number. Lateral and medial views are shown for the right (top) and left (bottom) hemispheres. The left panel shows the overall t-statistics for the relation in each parcel, while the right panel shows parcels where the relation between surface area and image quality survives multiple comparisons.

Figure S11.

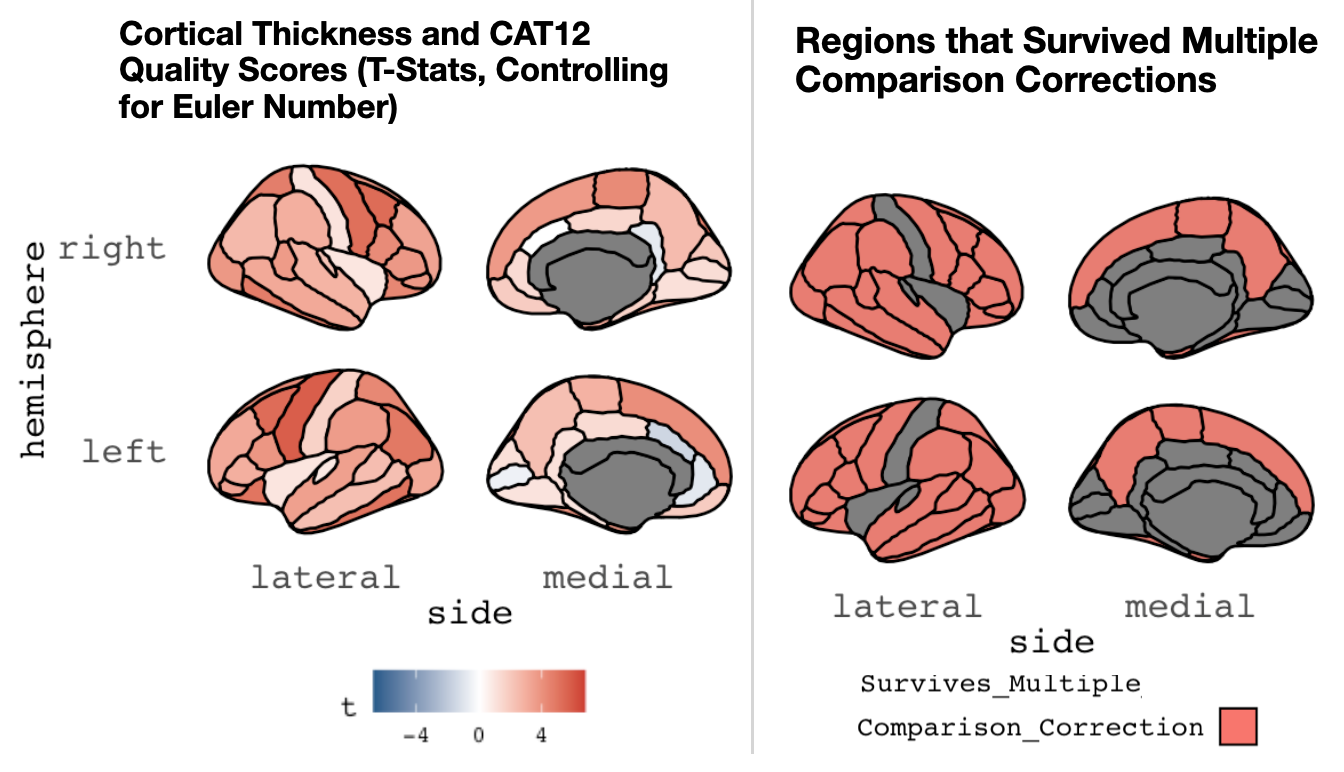

Caption: A graphic depiction (from the R library *ggseg*) showing associations between image quality (as assessed by the CAT12 toolbox) and derived (mean) cortical thickness. This is shown for the Desikan atlas commonly used in Freesurfer. These analyses controlled for Freesurfer Euler Number. Lateral and medial views are shown for the right (top) and left (bottom) hemispheres. The left panel shows the overall t-statistics for the relation in each parcel, while the right panel shows parcels where the relation between surface area and image quality survives multiple comparisons.

Figure S12.

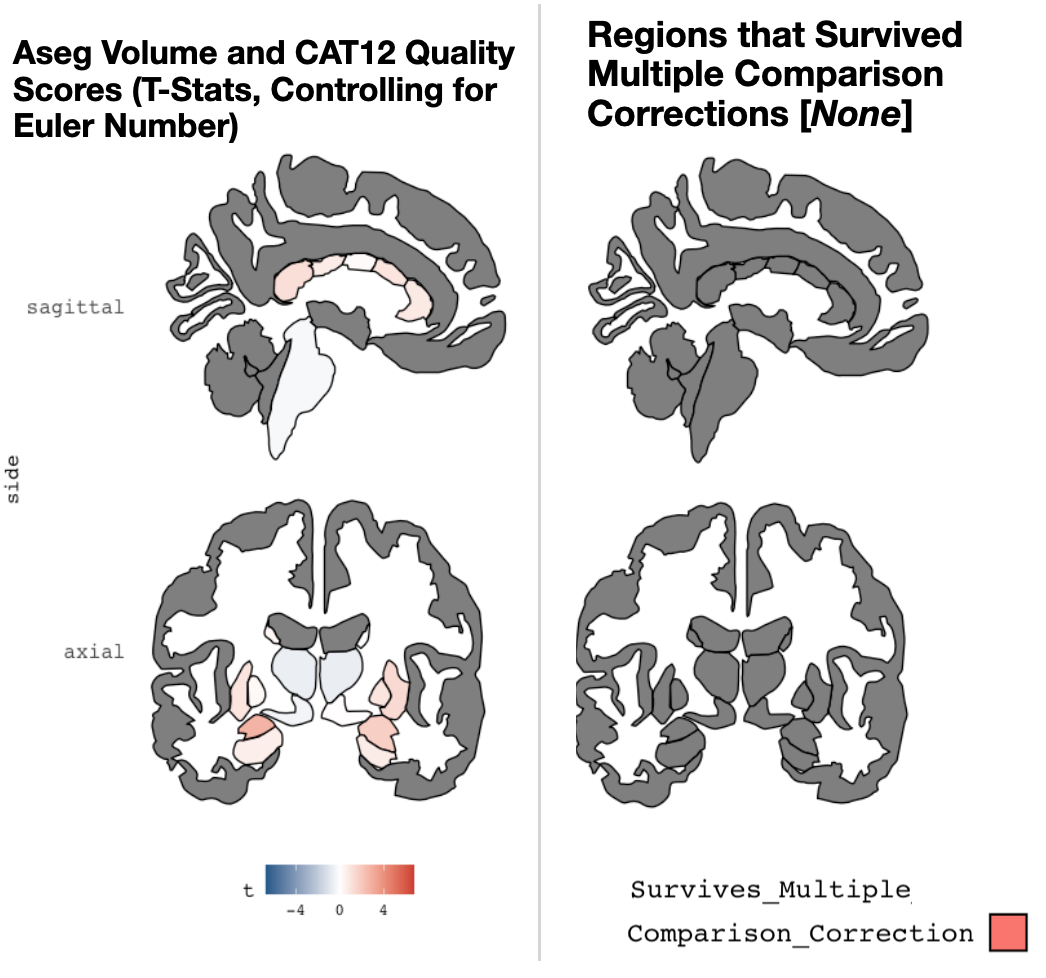

Caption: A graphic depiction (from the R library ggseg) showing associations between image quality (assessed by the CAT12 Toolbox) and subcortical volumes. This is shown for the Freesurfer ASEG atlas. Coronal (left) and sagittal (right) views are shown. These analyses controlled for Freesurfer’s Euler Number. The left panel shows the t-statistic for the relation in each subcortical volume, while the right panel shows parcels where the relation between volume and image quality survives multiple comparisons. Of note, no regions survived multiple comparisons correction.

Figure S13.

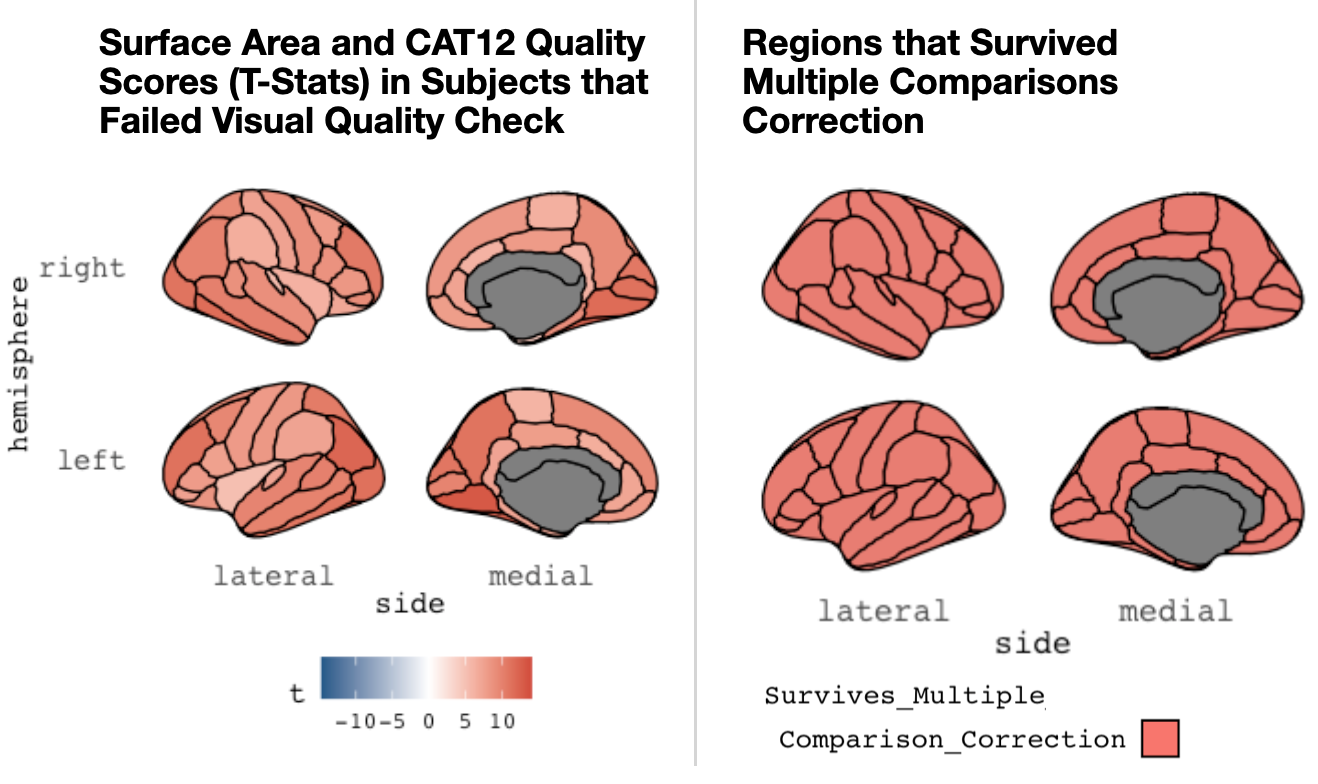

Caption: A graphic depiction (from the R library *ggseg*) showing associations between image quality (as assessed by the CAT12 toolbox) and derived (mean) cortical surface area. This is shown for the Desikan atlas commonly used in Freesurfer. These analyses were completed in participants who failed visual quality control checks (by human raters). Lateral and medial views are shown for the right (top) and left (bottom) hemispheres. The left panel shows the overall t-statistics for the relation in each parcel, while the right panel shows parcels where the relation between surface area and image quality survives multiple comparisons. Importantly, the color scale is changed compared to other similar graphics previously shown in this supplement.

Figure S14.

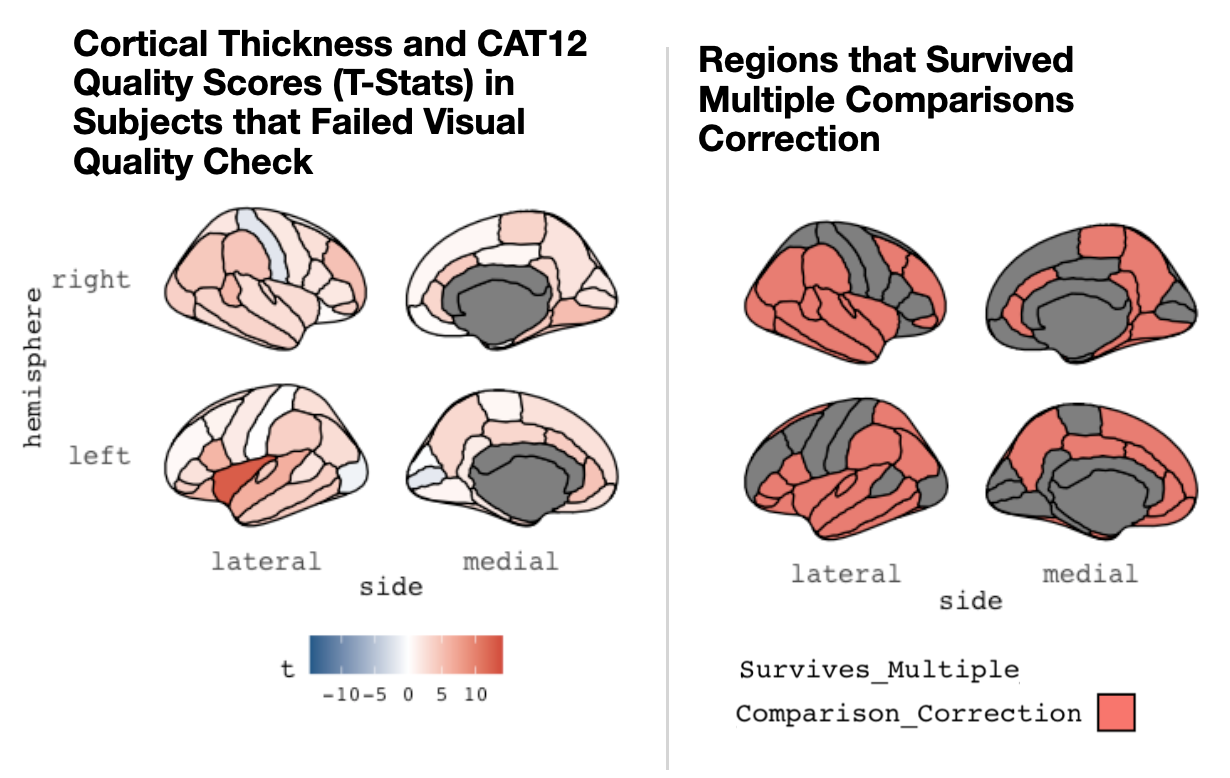

Caption: A graphic depiction (from the R library *ggseg*) showing associations between image quality (as assessed by the CAT12 toolbox) and derived (mean) cortical thickness. This is shown for the Desikan atlas commonly used in Freesurfer. These analyses were completed in participants who failed visual quality control checks (by human raters). Lateral and medial views are shown for the right (top) and left (bottom) hemispheres. The left panel shows the overall t-statistics for the relation in each parcel, while the right panel shows parcels where the relation between surface area and image quality survives multiple comparisons. Importantly, the color scale is changed compared to other similar graphics previously shown in this supplement.

Figure S15.

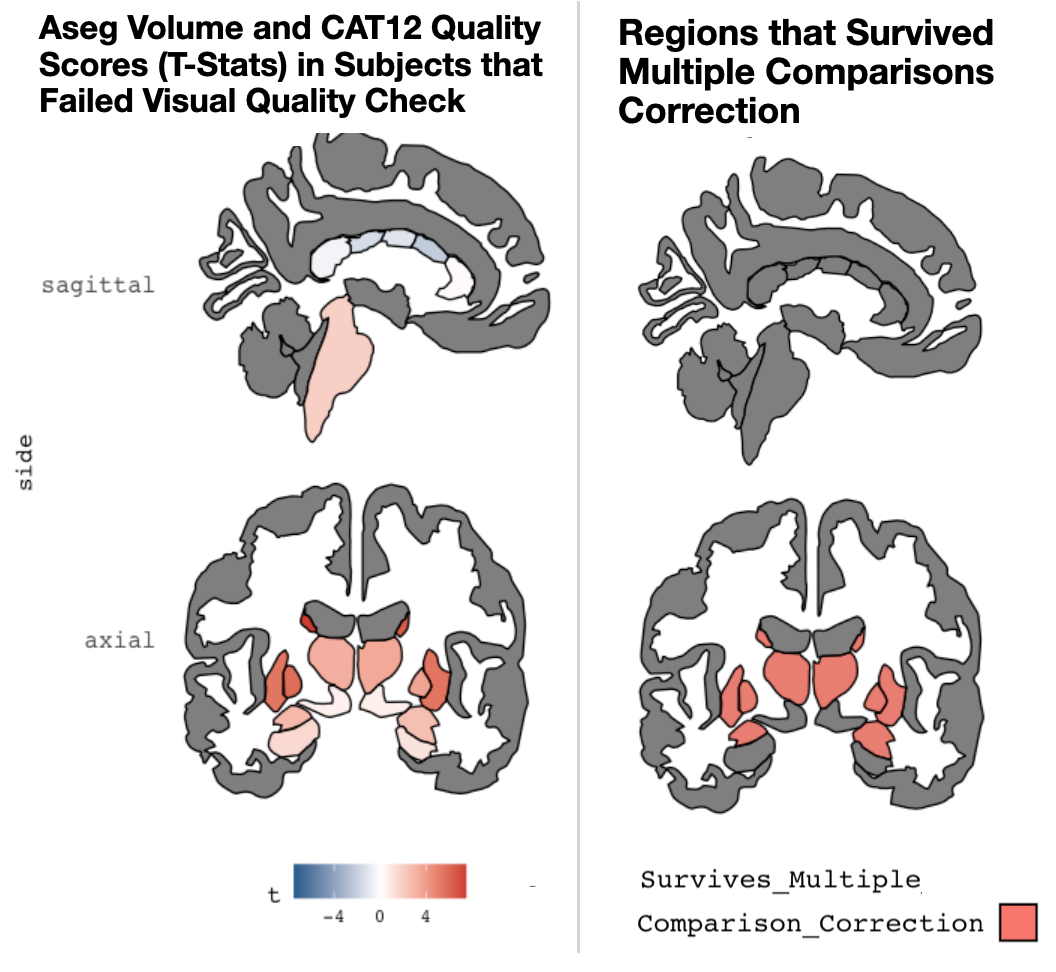

Caption: A graphic depiction (from the R library ggseg) showing associations between image quality (assessed by the CAT12 Toolbox) and subcortical volumes. This is shown for the Freesurfer ASEG atlas. These analyses were completed in participants who failed visual quality control checks (by human raters). Coronal (left) and sagittal (right) views are shown. The left panel shows the t-statistic for the relation in each subcortical volume, while the right panel shows parcels where the relation between volume and image quality survives multiple comparisons.

Figure S16.

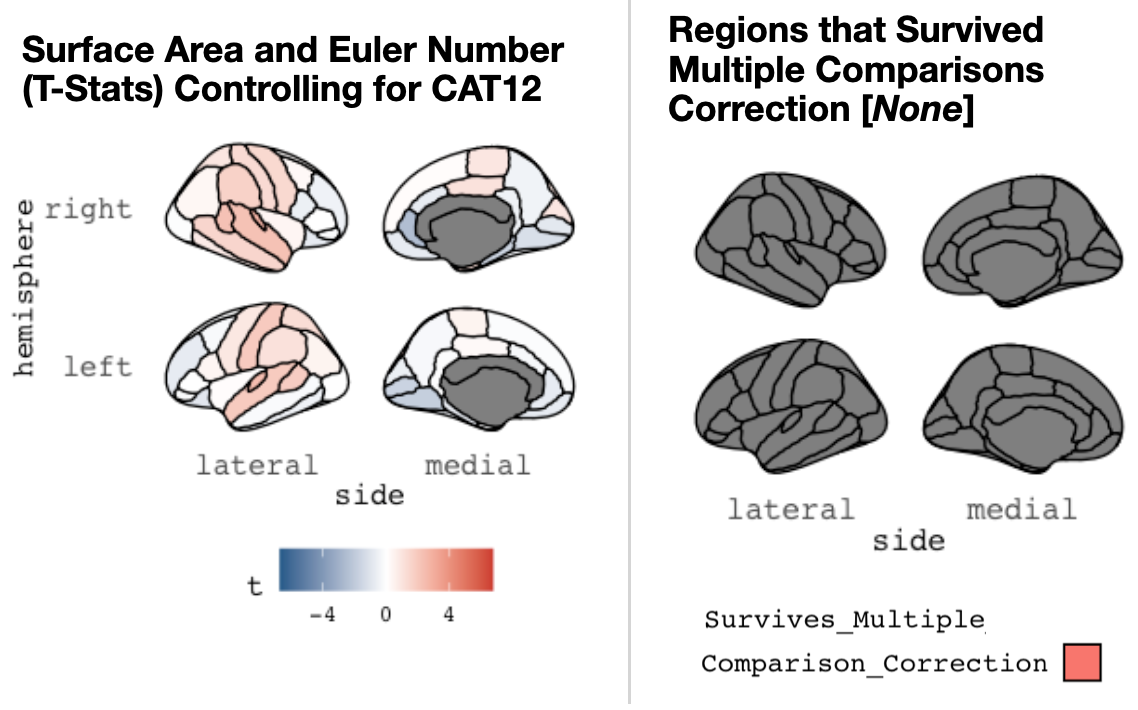

Caption: Caption: A graphic depiction (from the R library ggseg) showing associations between Euler Number and derived (mean) cortical surface area. These analyses control for CAT12 quality score. Results are shown for the Desikan atlas commonly used in Freesurfer. Lateral and medial views are shown for the right (top) and left (bottom) hemispheres. The left panel shows the overall t-statistics for the relation in each parcel, while the right panel shows parcels where the relation between surface area and image quality survives multiple comparisons. Of note, no regions survived multiple comparisons correction.

Figure S17.

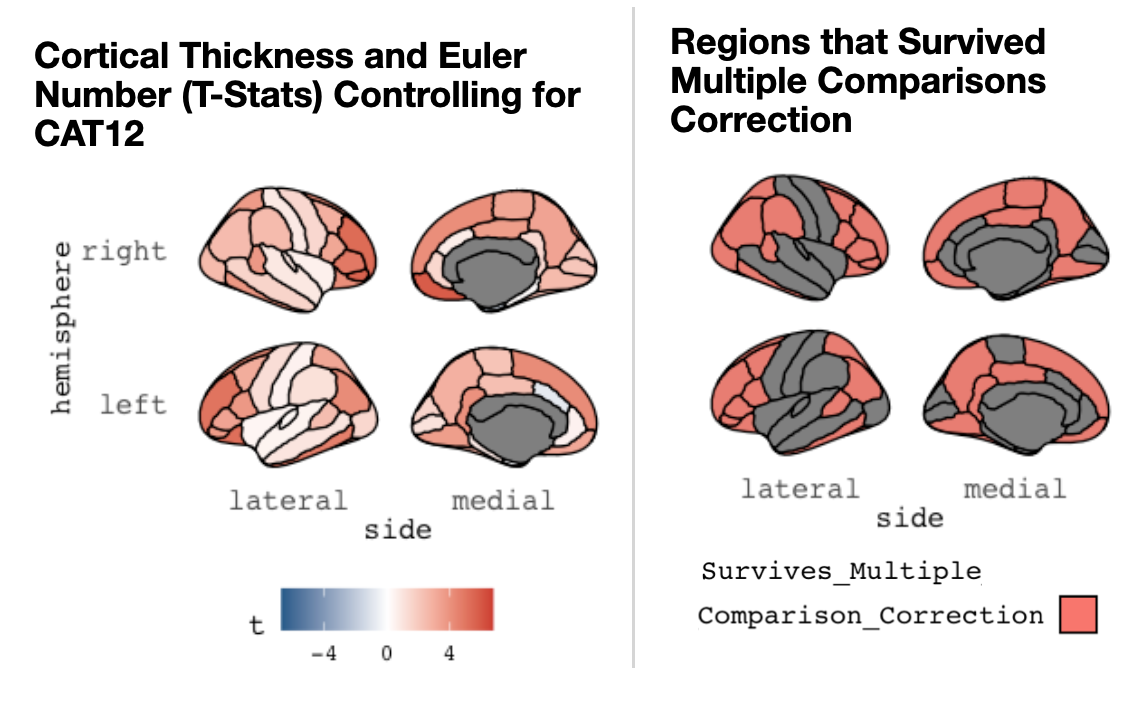

Caption: A graphic depiction (from the R library ggseg) showing associations between Euler Number and derived (mean) cortical thickness. These analyses control for CAT12 quality score. Results are shown for the Desikan atlas commonly used in Freesurfer. Lateral and medial views are shown for the right (top) and left (bottom) hemispheres. The left panel shows the overall t-statistics for the relation in each parcel, while the right panel shows parcels where the relation between surface area and image quality survives multiple comparisons.

Figure S18.

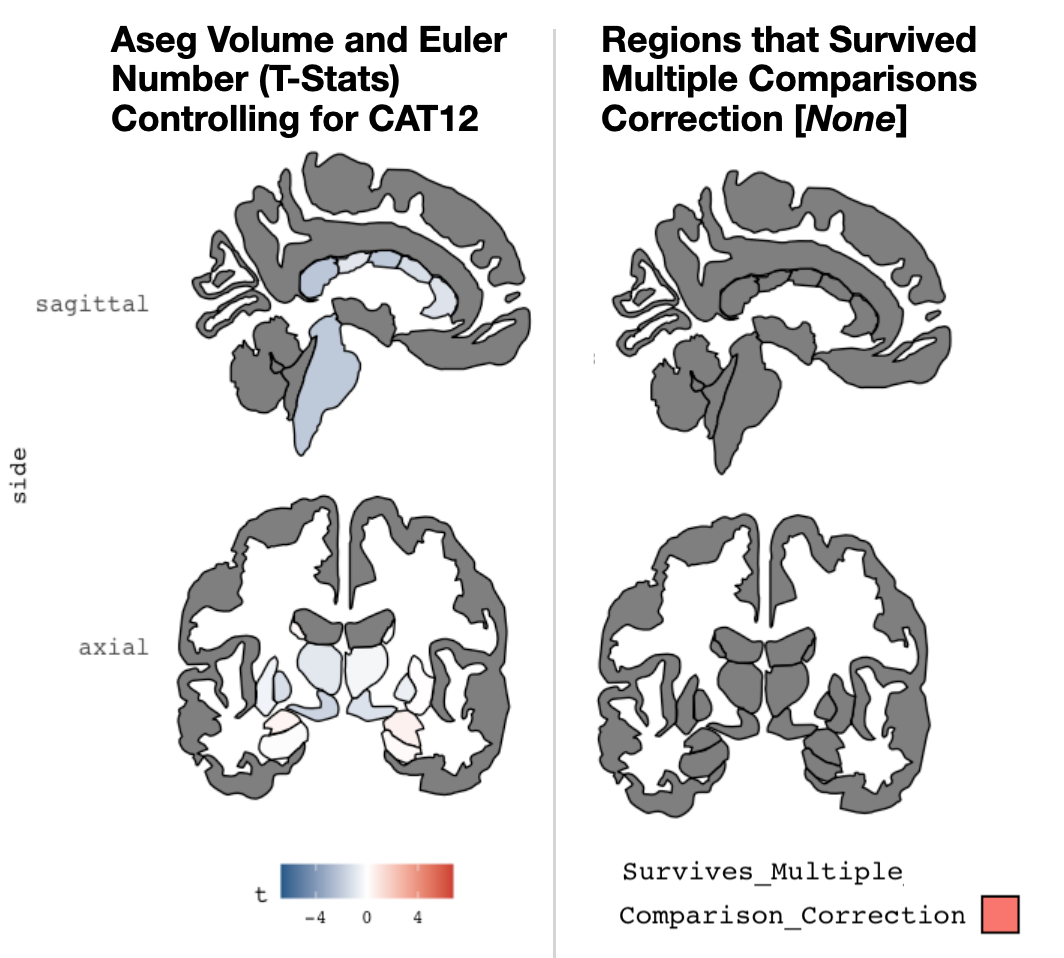

Caption: A graphic depiction (from the R library ggseg) showing associations between Euler Number and subcortical volumes. These analyses control for CAT12 quality score. This is shown for the Freesurfer ASEG atlas. Coronal (left) and sagittal (right) views are shown. The left panel shows the t-statistic for the relation in each subcortical volume, while the right panel shows parcels where the relation between volume and image quality survives multiple comparisons.
